# The *Legionella* effector RidL promotes mitochondrial fragmentation through phosphorylation activation of the large GTPase Drp1

**DOI:** 10.1101/2025.10.07.680855

**Authors:** Ana Katic, Elizabeth Teresa Vittori, Stefanie Halter, Kevin Steiner, A. Leoni Swart, Milad Radiom, Francisco Javier García-Rodríguez, Tobias Jäggi, Xiaodan Li, Jason A. Mears, Carmen Buchrieser, Pedro Escoll, Vikram Govind Panse, Hubert Hilbi

**Author notes:** Contributed equally (Ana Katic, Elizabeth Teresa Vittori). Institute of Medical Virology, University of Zürich; Zürich, Switzerland (Kevin Steiner). Biozentrum, University of Basel; Basel, Switzerland (A. Leoni Swart).

## Abstract

Intracellular pathogens such as *Legionella pneumophila* secrete effector proteins that manipulate host cell processes to promote bacterial survival. One such effector, RidL, is known to inhibit retrograde trafficking by interacting with the retromer complex via its N-terminal domain. Here, we identify a second function of RidL mediated by its C-terminal domain, which directly binds to the mitochondrial fission GTPase Drp1 and related large GTPases. In vitro, RidL reduces Drp1 GTPase activity and disrupts its oligomerization. During infection, RidL localizes to mitochondria, enhances the accumulation of Drp1 and the outer membrane protein Tom20, and impairs mitochondrial dynamics and function. Moreover, in *L. pneumophila*-infected cells, RidL promotes phosphorylation of Drp1 at Ser616, leading to Drp1 activation and mitochondrial fragmentation. These findings establish RidL as a bifunctional effector that targets both the retromer complex and Drp1 through distinct domains. By interfering with host mitochondrial dynamics, RidL enables *L. pneumophila* to remodel host organelles and optimize conditions for intracellular replication.

## Introduction

The environmental bacterium *Legionella pneumophila* is a facultative intracellular pathogen, which can cause a life-threatening pneumonia called Legionnaires’ disease ^1,2^. *L. pneumophila* manipulates eukaryotic host cells in a highly sophisticated manner and replicates intracellularly in a unique compartment termed the “*Legionella*-containing vacuole” (LCV) ^1,2^. To govern pathogen-host cell interactions, *L. pneumophila* employs the Icm/Dot type IV secretion system (T4SS) that translocates more than 300 different “effector proteins” into host cells, where they subvert pivotal processes, including vesicle trafficking, cytoskeletal dynamics, and signal transduction ^3,4^. Some of the secreted effectors target the mitochondria or their dynamics ^5^, and act as a nucleotide carrier protein ^6^, modify ADP/ATP translocases by reversible ADP-ribosylation ^7,8^, cleave syntaxin 17 ^9^, or stabilize microtubules as Ran GTPase activators ^10–12^.

Mitochondria are highly dynamic organelles, and the fission, fusion, trafficking and interactions of these organelles are essential for their function as cellular “powerhouses” and signaling hubs, controlling metabolism, proliferation, and apoptosis ^13,14^. Defects in these processes are linked to ageing ^15^ and many diseases, including neurodegenerative disorders ^16^, cardiovascular ailments ^17^, and cancer ^18^.

A pivotal regulator of mitochondrial dynamics is the 78 kDa large fission GTPase dynamin-related protein 1 (Drp1), which acts in several steps: (i) subcellular translocation from the cytoplasm to the outer mitochondrial membrane, (ii) assembly into multimeric tubular structures called spirals, and (iii) GTP hydrolysis associated with membrane “constriction” followed by disassembly ^14^. Drp1 activity is regulated by posttranslational modifications, including activation by phosphorylation at Ser616 ^19^. Another regulator of mitochondrial dynamics is the retromer coat complex, comprising the cargo recognition subcomplex subunits Vps35, Vps29, and Vps26, as well as membrane-deforming sorting nexins (SNXs) ^16,20^.

The retromer complex and Drp1 regulate the formation of mitochondria-derived vesicles (MDVs), operating in mitochondrial quality control and protein turnover pathways that transport selected cargo proteins and lipids to lysosomes or peroxisomes for degradation ^21–25^. In some MDV pathways, the retromer initiates membrane curvature, followed by the elongation of membrane protrusions pulled along microtubule filaments and scission by Drp1. Specifically, the retromer Vps35 subunit regulates the MDV-dependent turnover of Drp1 itself ^26,27^ as well as the mitochondrial outer membrane protein Tom20, a component of the import receptor complex transporting cytoplasmic mitochondrial precursor proteins ^24,25^. The fusion of MDVs with lysosomes is mediated by the SNARE syntaxin 17 ^28^.

An imbalance in retromer- and Drp1-dependent mitochondrial dynamics underlies neurodegenerative disorders. In Huntington’s disease, mutant forms of the protein huntingtin abnormally interact with Drp1 and stimulate its enzymatic activity ^29^. In Parkinson’s disease, mutant forms of Vps35 showed an enhanced interaction with Drp1, which promotes the removal of inactive Drp1 complexes via MDV-dependent trafficking to lysosomes, leading to the polymerization of new active Drp1 and mitochondrial fragmentation ^26,27^.

*L. pneumophila* produces a 131 kDa effector protein called RidL (retromer interactor decorating LCVs), which promotes intracellular replication, inhibits retrograde transport, and binds the Vps29 subunit of the retromer cargo recognition subcomplex ^30^. The interaction of RidL with Vps29 occurs through a small amino acid stretch in the effector’s N-terminal fragment termed β-loop, and leads to the displacement on Vps29 of the Rab7 GTPase activating protein (GAP) TBC1D5 ^30–33^. The interactor(s) and function of the C-terminal fragment of RidL are unknown. In this study, we identify the large mitochondrial GTPase Drp1 as an interactor of the C-terminal fragment of RidL, and we assess the role of RidL for the phosphorylation activation of Drp1, and its consequences for mitochondrial dynamics and functions.

## Results

### RidL binds to Drp1 and impairs GTPase activity

Previous structural studies of RidL revealed an N-terminal “foot-like” fold that at its “heel” binds via the “β-loop” to the Vps29 retromer subunit ^31–33^. The experimental results closely match the Alphafold2 structure prediction (**Fig. 1A**), which further stipulates a “leg” domain, which is linked by an apparently flexible hinge to domains resembling an “arm” and a “fist”. According to the model, the C-terminus of RidL folds back along the “arm” domain as an extended, kinked α-helix and ends in the hinge region. The function and potential interaction partners of the C-terminal fragment of RidL are unknown. Based on these structural elements of RidL, we sought to identify further interaction partners of the effector.

**Fig. 1.**
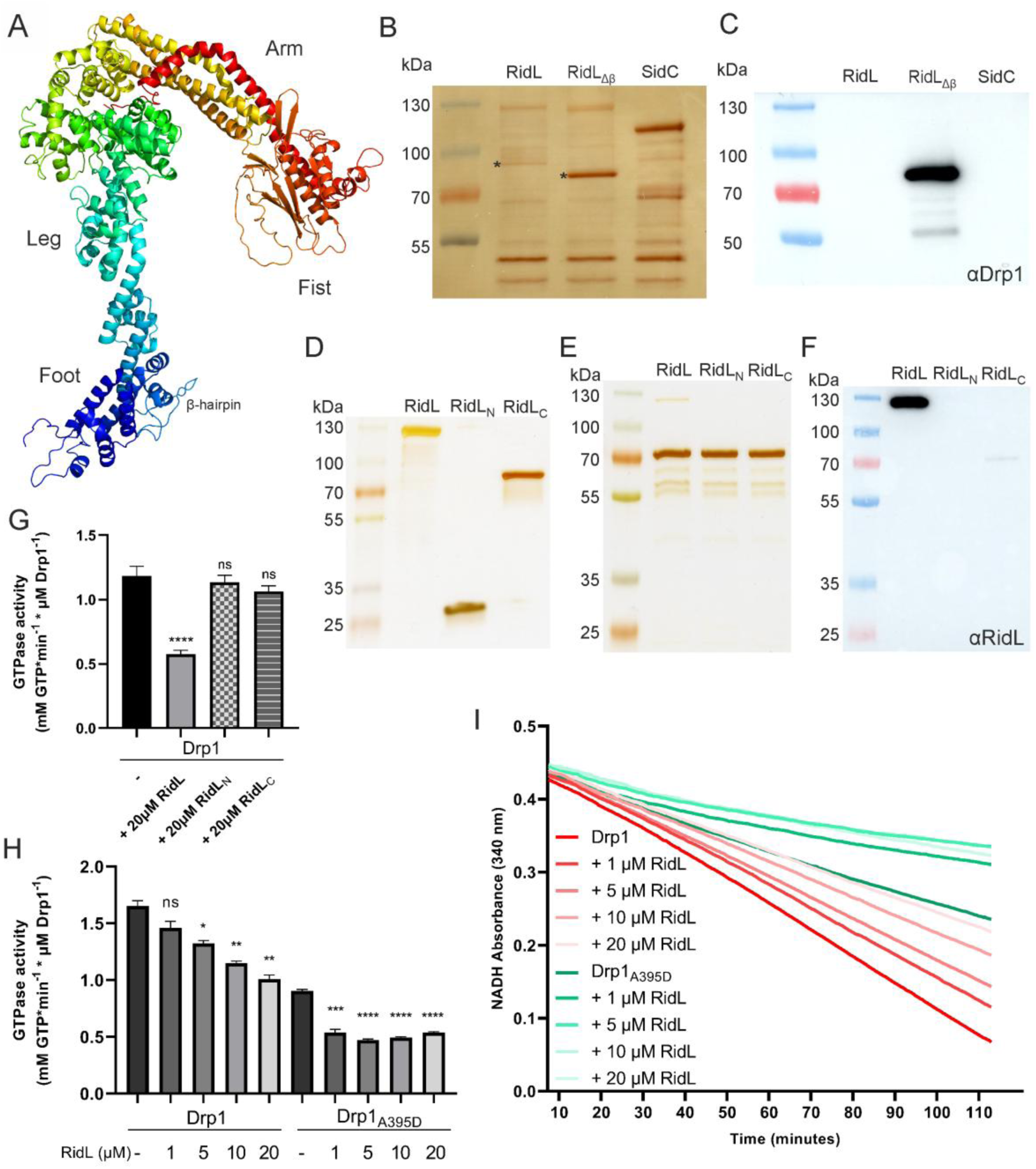
RidL binds to Drp1 and impairs GTPase activity *in vitro*. (**A**) Alphafold2 model of RidL. The model shows the “foot”, “leg”, “arm”, and “fist” domains. (**B**, **C**) Purified RidL, RidL_Δβ_ (lacking the Vps29-binding β-loop), or SidC was coupled to UltraLink Biosupport beads, incubated with HeLa cell lysate, and washed once. Proteins bound to washed beads were separated by SDS-PAGE and analyzed by (**B**) silver stain and mass spectrometry (protein bands labelled by stars), or (**C**) anti-Drp1 Western blot. (**D**) Purified RidL (131 kDa), RidL_N_ (RidL_1-258_, 30 kDa), or RidL_C_ (RidL_437-1100_, 75 kDa) were visualized by silver-stain, and equal amounts were incubated with purified Drp1 (78 kDa) coupled to UltraLink Biosupport beads. The beads were washed once, and bound proteins were separated by SDS-PAGE and visualized by (**E**) silver stain, or (**F**) anti-RidL Western blot. (**G, H**) Coupled enzymatic assay to assess GTPase activity of 1 µM purified Drp1 or the oligomerization mutant Drp1_A395D_ in presence or absence of purified RidL, RidL_N_ (RidL_1-258_), or RidL_C_ (RidL_437-1100_): Data from continuous NADH depletion assays converted to Drp1 GTPase activity (2 technical duplicates each from 3 independent experiments, means + SEM; *, p < 0.05; **, p < 0.01; ***, p < 0.001; ****, p < 0.0001). (**I**) Graphical depiction of continuous NADH depletion over time reflecting Drp1 GTPase activity (means of 2 technical duplicates each from 3 independent experiments).

To assess RidL binding partners in lysates of mammalian cells, we performed pulldown experiments with purified RidL, RidL_Δβ_ (lacking the Vps29-binding β-loop, **Fig. 1A**) and – as a negative control – the *L. pneumophila* effector SidC. Beads coupled with these baits were incubated with HeLa cell lysate and washed. Bound proteins were eluted, separated by SDS-PAGE, and dominant bands were identified by mass spectrometry (MS) (**Fig. 1B**). Thus, the retromer coat complex subunit Vps35 was identified as an (indirect) binding partner of RidL as observed previously ^30^, and – intriguingly – the large fission GTPase Drp1 was identified as a potential interaction partner of RidL_Δβ_. The binding of Drp1 in HeLa cell lysates to RidL_Δβ_ was validated by Western blot (**Fig. 1C**). Taken together, full-length RidL binds to the retromer complex in HeLa cell lysates, and RidL_Δβ_ (no longer interacting with the retromer) binds to Drp1.

To test whether and which portion of RidL directly binds to Drp1 *in vitro*, purified Drp1 was immobilized on beads and incubated with equal amounts of purified RidL, the N-terminal fragment RidL_N_ (RidL_1-258_), or the C-terminal fragment RidL_C_ (RidL_437-1100_) (**Fig. 1D**). The washed beads were subjected to SDS-PAGE, and bound proteins were visualized by silver stain (**Fig. 1E**) or anti-RidL Western blot (**Figs. 1F, S1A**). Using these purified proteins, full-length RidL strongly bound to Drp1, RidL_C_ bound to some extent, and RidL_N_ did not bind at all to Drp1. In summary, purified Drp1 binds *in vitro* to a C-terminal fragment, RidL_C_ (RidL_437-1100_), of RidL.

To determine whether RidL affects the GTPase activity of Drp1, we used a coupled GTPase enzyme assay ^34^ (**Fig. S1B**). Purified full-length RidL, RidL_N_ (RidL_1-258_) or RidL_C_ (RidL_437-1100_) was added to purified Drp1, and the GTPase activity was assessed (**Fig. 1G**). At 20× excess, RidL but neither RidL_N_ (RidL_1-258_) nor RidL_C_ (RidL_437-1100_) inhibited the GTPase activity. Upon adding increasing concentrations of purified RidL to purified Drp1 or to an assembly-deficient mutant, Drp1_A395D_ ^35^, RidL inhibited the GTPase activity of wild-type Drp1 in a dose-dependent but inefficient manner (**Fig. 1H, I**): at 20× excess, the activity was reduced less than two-fold. RidL also reduced the GTPase activity of Drp1_A395D_ by approximately two-fold, indicating that the inhibitory effect was independent of GTPase oligomerization. Taken together, these results indicate that an excess of full length RidL but not fragments thereof inhibit the GTPase activity of Drp1 and an oligomerization-deficient mutant protein.

### RidL impairs Drp1 oligomerization and spiralization

Drp1 is a large fission GTPase, which – with complex dynamics – oligomerizes and forms spirals around membranes to constrict and sever membrane bilayers, leading to mitochondrial fragmentation as well as the release of MDVs ^24,36^. To assess effects of RidL on the oligomerization and spiralization of Drp1, we employed negative-staining electron microscopy (EM) (**Fig. 2A**). Using this approach, we sought to validate that upon treatment of purified Drp1 with the non-hydrolysable GTP analogue guanosine-5’-[(β,γ)-methyleno]triphosphate (GMPPCP), Drp1 oligomerizes and forms spirals ^37,38^. Under the conditions used (0.5 mM GMPPCP), the majority of GMPPCP-treated Drp1 indeed oligomerized and formed spirals, which sometimes clustered and aligned in parallel. Intriguingly, the concomitant addition of excess RidL disrupted oligomerization and coiling of GMPPCP-treated Drp1, while the addition of purified SidC, an unrelated *L. pneumophila* effector of similar size, did not (**Fig. 2A**). Thus, RidL specifically impairs the oligomerization and spiralization of Drp1, which might contribute to the weak inhibition of Drp1 GTPase activity observed (**Fig. 1G-I**).

**Fig. 2.**
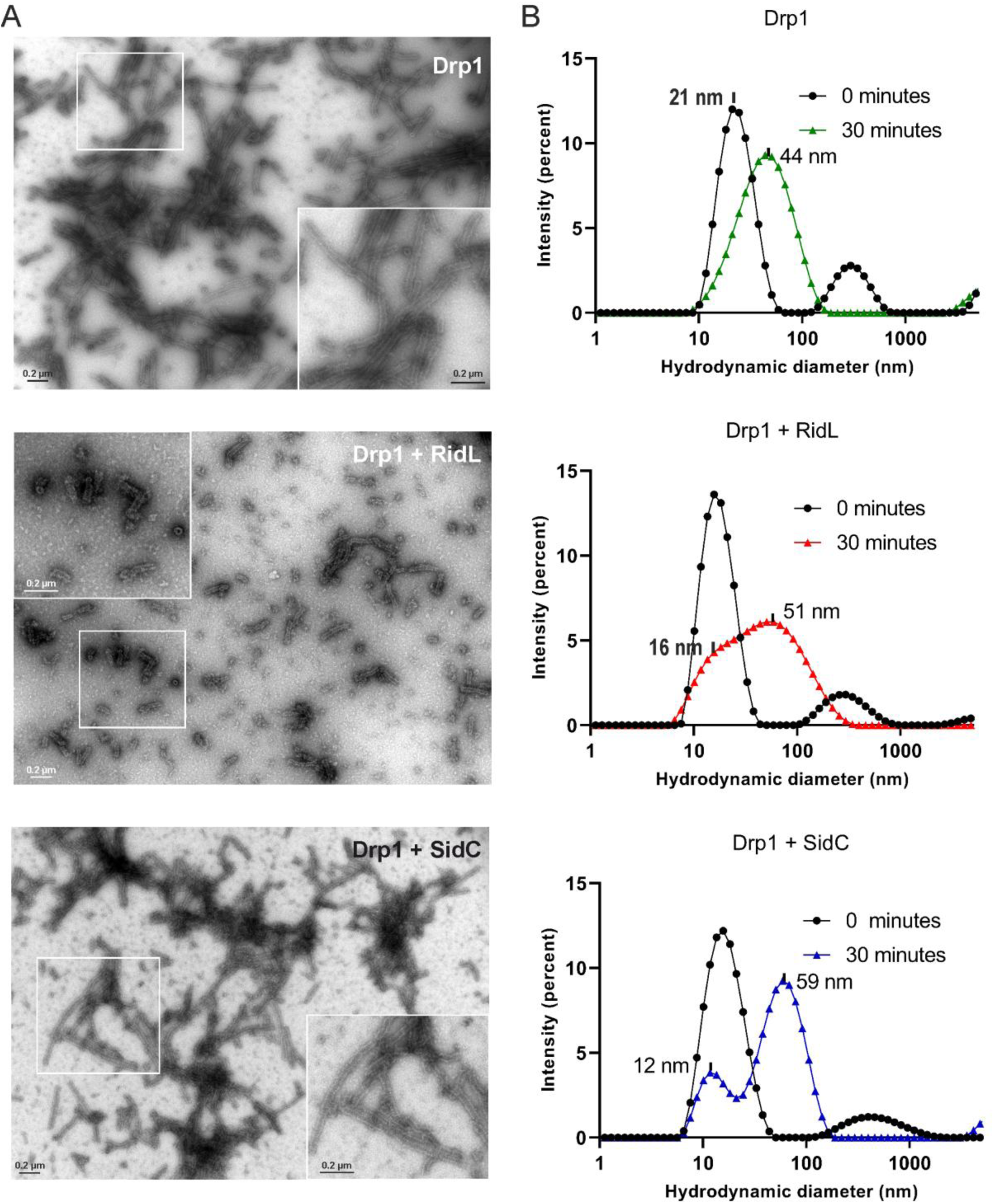
RidL inhibits Drp1 oligomerization and tubulation. (**A**) Purified Drp1 (5 µM) was treated with 0.5 mM GMPPCP, and RidL (15 µM), SidC (15 µM), or left untreated, and incubated for 1 h (room temperature) before staining. Oligomerization/spiralization was assessed by negative-stain electron microscopy. Scale bars, 0.2 µm; inset 0.2 µm. (**B**) Purified Drp1 (1 µM) was incubated with 0.5 mM GMPPCP and treated with RidL (3 µM), SidC (3 µM), or left untreated, and Drp1 oligomerization was assessed by dynamic light scattering.

Analogously, we assessed the effect of RidL (131 kDa) on the oligomerization of Drp1 (78 kDa) by dynamic light scattering (DLS) (**Fig. S2**). This approach indicated that upon treatment of purified Drp1 with 0.5 mM GMPPCP, the hydrodynamic diameter of Drp1 complexes increased (**Fig. 2B**). Prior to the addition of GMPPCP (0 min), the peak around 21 nm likely corresponds to the primarily dimeric form of Drp1 (156 kDa) and multiples thereof ^39^. Within 30 min of GMPPCP treatment, a complex oligomeric Drp1 population formed, which showed a broader distribution peaking at a higher hydrodynamic diameter around 44 nm. Upon addition of excess RidL to GMPPCR-treated Drp1, a broad population of complexes formed within 30 min, including the peak around 16 nm associated with RidL (**Figs. 2B**, **S2B**) and showing a peak at 51 nm. Based on the finding that RidL binds Drp1 (**Fig. 1BC**), the peak around 51 nm likely shows a population of aggregated RidL-Drp1 complexes. Finally, upon addition of excess SidC (103 kDa), a size distribution with two peaks was observed within 30 min, with the first peak at 12 nm associated with SidC (**Fig. 2B**, **Fig. S2C**) and a second peak around 59 nm. Based on the finding that SidC does not bind Drp1 (**Fig. 1BC**), the second peak around 59 nm likely corresponds to oligomerized Drp1, which might form even larger aggregates than in absence of SidC. In summary, negative-stain EM and DLS revealed that RidL but not SidC impairs oligomerization and spiralization of GMPPCP-treated Drp1.

### RidL interacts with evolutionary conserved large fission GTPases

RidL and other *L. pneumophila* effectors function in evolutionary distant host cells and subvert conserved targets ^30,31,40^. To validate the binding of RidL to Drp1 and possibly identify additional eukaryotic large fission GTPases as target(s) of RidL, we employed yeast two-hybrid (Y2H) assays using RidL as the bait and large fission GTPases from mammalian cells, *Dictyostelium discoideum* amoeba and the yeast *Saccharomyces cerevisiae* as the prey (**Fig. 3A**). Using this approach, RidL was found to interact with large fission GTPases from mammalian cells (Drp1 and Dnm1, but not Dnm2 or Dnm3), *D. discoideum* (DymA, but not DymB) and *S. cerevisiae* (Vps1). Interestingly, among the family of dynamin-like large fission GTPases, Drp1, DymA and Vps1 are the evolutionary most closely related enzymes ^41^.

**Fig. 3.**
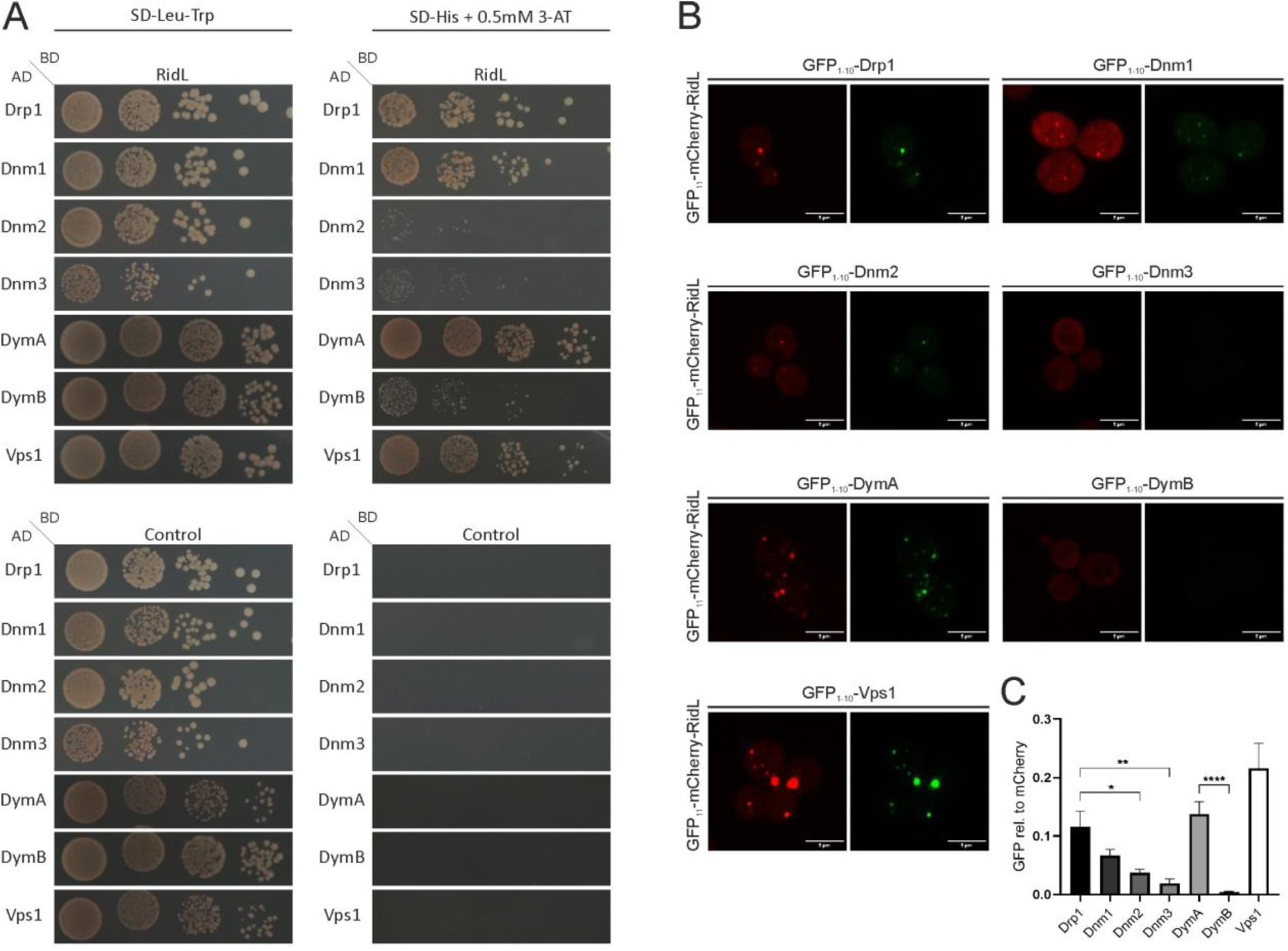
RidL interacts with evolutionary conserved large fission GTPases. (**A**) Yeast two-hybrid assays using the reporter strain NMY32 producing *L. pneumophila* RidL fused to the LexA DNA binding domain (BD) and large fission GTPases form human, *Dictyostelium discoideum* and yeast fused to the Gal4 activation domain (AD) of a split transcription factor required for His3 expression (negative controls: MTR4 fragment, lamin C). Yeast transformants were spotted in 10-fold serial dilutions on SD plates lacking Leu and Trp (SD-Leu-Trp) or on SD-His plates with 0.5 mM 3-AT (3 d, 30°C). Images are representative for 3 independent experiments. (**B**) Bimolecular fluorescence complementation (BiFC) in yeast strain BY4741 producing RidL fused to GFP_11_-mCherry, and large fission GTPases fused to GFP_1-10_. Aliquots of yeast transformants grown in selective medium were embedded in 0.1% agarose and imaged by confocal laser scanning microscopy. (**C**) Quantification of (B). Integrated densities of GFP and mCherry signals in merged z-stacks were calculated by Image J, and GFP signal intensities (representing RidL-GTPases interaction) were normalized to mCherry intensities representing overall protein production levels. Graphs show means + SEM of relative GFP intensities of nine yeast transformants from three independent experiments.

As an alternative approach, we tested the interaction of RidL with large fission GTPases using confocal laser scanning microscopy and bimolecular fluorescence complementation (BiFC) in yeast. To this end, RidL was fused to GFP_11_-mCherry, and large fission GTPases were fused to GFP_1-10_ (**Fig. 3B**). RidL interacted and formed GFP-positive, punctate structures in the cells with Drp1 (and to a lesser extent with Dnm1), DymA and Vps1. Quantification of the interactions revealed the following intensity: Vps1 > DymA > Drp1 > Dnm1 > Dnm2 >

Dnm3 > DymB, where the latter three fission GTPases did not (or only very weakly) interact with RidL (**Fig. 3C**). In summary, RidL interacts most strongly with the closely related large fission GTPases Drp1, DymA, Vps1, as well as with Dnm1.

### RidL is associated with mitochondria and increases mitochondrial Drp1 and Tom20

To functionally validate the Y2H and BiFC interaction experiments, we assessed whether Drp1 or Dnm1 are implicated in intracellular replication of *L. pneumophila*. To this end, Drp1 or Dnm1 were depleted by RNA interference (RNAi) in A549 epithelial cells, and intracellular bacterial replication was assessed (**Fig. 4A**). Whereas the depletion of Drp1 impaired the intracellular replication of *L. pneumophila* as previously observed ^11^, the depletion of Dnm1 had no effect. RNAi efficiently depleted the target proteins Drp1 and Dnm1 (**Fig. S3AB**) and was not cytotoxic for the cells (**Fig. S3CD)**. Taken together, while RidL interacts both with Drp1 and Dnm1, only Drp1 but not Dnm1 promotes the intracellular growth of *L. pneumophila*. Given that Drp1 localizes to and is implicated in mitochondrial dynamics, we next checked whether RidL is associated with mitochondria in *L. pneumophila*-infected cells. To this end, HeLa cells were infected with the *L. pneumophila* parental strain JR32, the Δ*icmT* mutant strain lacking a functional T4SS, the Δ*ridL* mutant strain lacking RidL, or the complemented Δ*ridL* mutant. Mitochondria were isolated by differential centrifugation and sucrose density gradient ultracentrifugation, and RidL was detected by Western blot in cytoplasmic (cyt) and purified mitochondrial (mito) fractions (**Fig. 4B**). Upon infection with *L. pneumophila*, RidL was detected in the mitochondrial fractions isolated from the parental strain JR32 and the complemented *ΔridL* mutant strain, but not from the mitochondrial fractions of cells infected with the *ΔicmT* or *ΔridL* mutant strains. While RidL was only detectable in the mitochondrial fraction in cells infected with strain JR32, the effector localized to the mitochondrial as well as the cytoplasmic fraction upon overproduction in the complemented *ΔridL* mutant strain (**Fig. 4B**). These results indicate that RidL localizes to mitochondria in *L. pneumophila*-infected cells, and a functional Icm/Dot T4SS is required for subcellular localization of the effector.

**Fig. 4.**
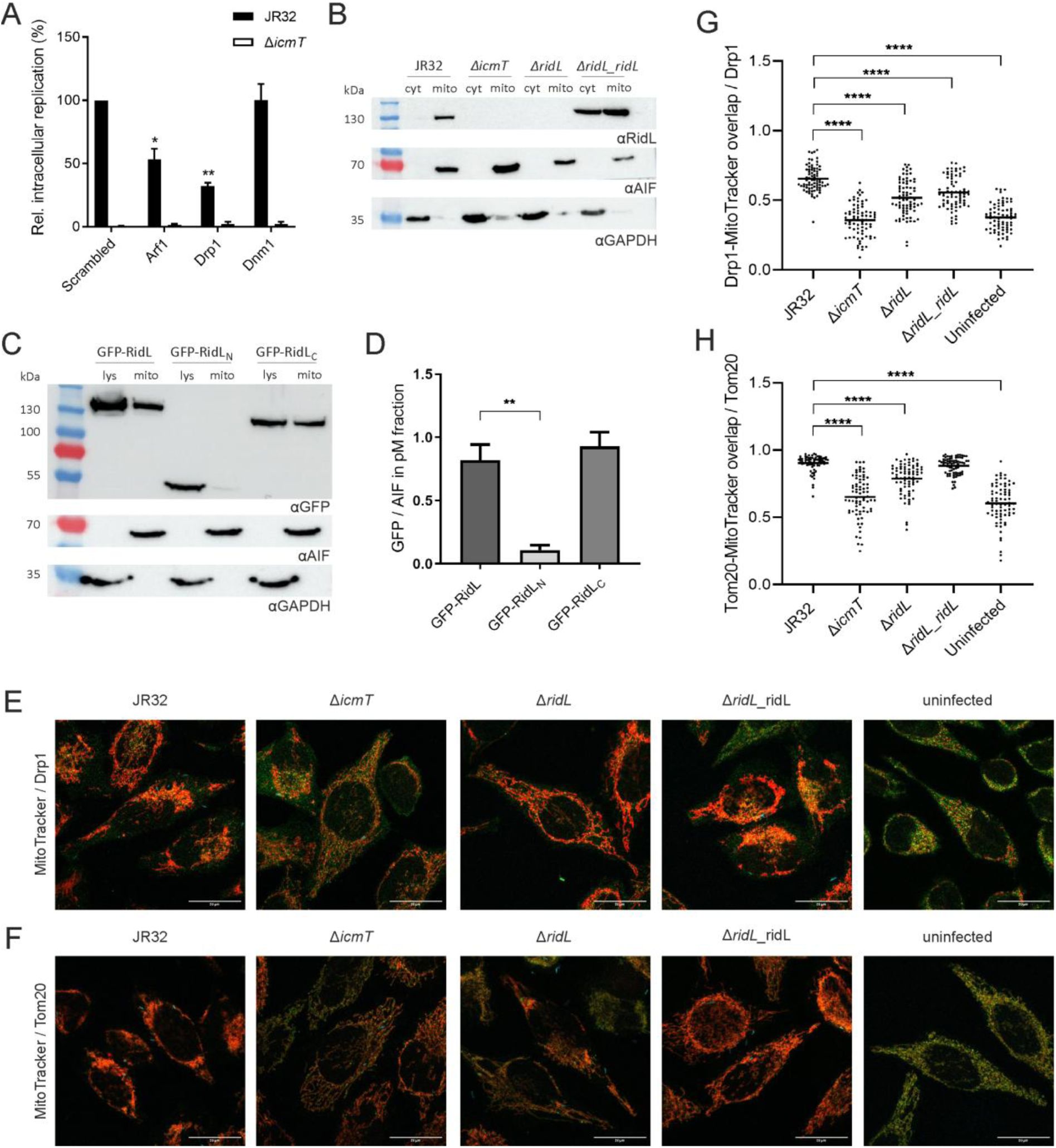
RidL is associated with mitochondria and increases mitochondrial Drp1 and Tom20. (**A**) A549 cells depleted (48 h) by siRNA for Arf, Drp1, or Dnm1 were infected (MOI 10, 24 h) with GFP-producing *L. pneumophila* JR32 or Δ*icmT* (pNT28), and intracellular replication was assessed by GFP fluorescence using a microtiter plate reader. Graphs show the relative intracellular replication (normalized to GFP signal 1 h p.i.) and represent means + SEM of three independent experiments (Student’s *t*-test, *, p < 0.05; **, p < 0.01). (**B**) Mitochondria were isolated by differential centrifugation and sucrose density gradient ultracentrifugation from HeLa cells infected (MOI 100, 6 h) with GFP-producing *L. pneumophila* JR32, *ΔicmT*, *ΔridL*, or *ΔridL*_*ridL* (pNT28 or pIF009). Western blot of RidL in cytoplasmic (cyt) and purified mitochondrial (mito) fractions; apoptosis-inducing factor (AIF) and glyceraldehyde 3-phosphate dehydrogenase (GAPDH) served as a mitochondrial or cytoplasmic marker, respectively. The figure shown is representative for 3 independent experiments. (**C**) HEK293 cells were transfected (24 h) for ectopic production of GFP-RidL (RidL, pKB248), GFP-RidL_9-258_ (RidL_N_, pKB249) or GFP-RidL_259-1167_ (RidL_C_, pKB250), mitochondria were isolated, and the GFP fusion proteins were detected by anti-GFP Western blot in the cell lysate (lys) and purified mitochondrial (mito) fractions. The figure shown is representative for 3 independent experiments. (**D**) Quantification of (C). Signal intensities of GFP and AIF were calculated by Image J. Bars show relative GFP signal in pure mitochondria fractions. Means + SEM of 3 independent experiments are shown (Student’s *t-*test, **, p < 0.01). (**E, F**) Fluorescence micrographs of HeLa cells infected (MOI 25, 2 h) with mCerulean-producing *L. pneumophila* JR23, Δ*icmT*, Δ*ridL* or Δ*ridL_ridL* (pNP99 or pKB208). Mitochondria were visualized with MitoTracker Deep Red, and immuno-labelled with specific antibodies against (**E**) Drp1 or (**F**) Tom20 (green) to quantify the localization of these proteins to mitochondria. (**G**) Quantification of signal overlap in (E) and (**H**) quantification of signal overlap in (F) show 75 infected cells from three independent experiments (each dot is a cell; Student’s *t*-test, ****, p < 0.0001). The brightness of fluorescence signals was linearly increased in Image J for enhanced visibility (A, B); original signal intensities of the images were processed for quantification (C, D).

To determine the portion of RidL that mediates its association to mitochondria, we transfected HEK293 cells with constructs for ectopic production of GFP-RidL, GFP-RidL_9-258_ (N-terminal fragment), or GFP-RidL_259-1167_ (C-terminal fragment) (**Fig. 1A**). Mitochondria were isolated from the transfected cells, and the GFP fusion proteins were detected by anti-GFP Western blots in the cell lysate (lys) and purified mitochondrial (mito) fractions (**Fig. 4C**). Under these conditions, GFP-RidL and GFP-RidL_259-1167_ but not GFP-RidL_9-258_ were associated with purified mitochondria (**Fig. 4D**), while all fragments were detected in similar amounts in the cell lysates (**Fig. S4A**). Accordingly, the C-terminal, Drp1-binding fragment RidL_259-1167_ determines the mitochondrial localization of the effector. In contrast, the N-terminal fragment RidL_9-258_, which binds the Vps29 subunit of the retromer cargo recognition subcomplex ^31^, does not determine the mitochondrial localization, even though Vps29 is present on mitochondria (**Fig. S4B**). In summary, our findings indicate that the retromer-binding effector RidL localizes through a C-terminal fragment to mitochondria, likely through its interaction with Drp1.

The retromer complex and the large fission GTPase Drp1 localize to mitochondria and promote MDV formation ^25^. Given that Drp1 and Tom20 are MDV cargo, we tested whether RidL affects the levels of these mitochondrial factors. To this end, HeLa cells were infected with the *L. pneumophila* parental strain JR32, Δ*icmT*, Δ*ridL*, or complemented Δ*ridL* mutant bacteria, and the mitochondrial levels of Drp1 (**Fig. 4E**, **Fig. S5A**) or Tom20 (**Fig. 4F**, **Fig. S5B**) were assessed by immuno-fluorescence microscopy. This approach revealed that upon presence and secretion of RidL, significantly more Drp1 (**Fig. 4G**) or Tom20 (**Fig. 4H**) localized to mitochondria, and the lower abundance of Tom20 upon infection with the Δ*ridL* mutant was fully reverted upon complementation of the mutant strain. Even less Drp1 or Tom20 localized to mitochondria upon infection of the cells with Δ*icmT* mutant bacteria. Taken together, in *L. pneumophila*-infected cells, RidL and a functional Icm/Dot T4SS increase the amounts of Drp1 and Tom20 on mitochondria.

### During *L. pneumophila* infection RidL promotes fragmentation and impairs function of mitochondria

The large fission GTPase Drp1 is a master regulator of mitochondrial dynamics ^14,19^. Given the localization of RidL to mitochondria and its role for mitochondrial Drp1 levels, we sought to assess whether the effector modulates mitochondrial dynamics and function. To this end, we stained HeLa cells with MitoTracker Orange, infected the cells with the parental strain JR32, Δ*icmT*, Δ*ridL*, or the complemented Δ*ridL* mutant strain, and visualized mitochondrial morphology by confocal microscopy (**Fig. 5A**). Compared to the *L. pneumophila* parental strain JR32, the mitochondrial aspect ratio (**Fig. 5B**) and branch length (**Fig. S6A**) were significantly increased upon infection with the Δ*ridL* strain, in particular at early time points post infection, and the effect was reversed upon infection with the complemented Δ*ridL* strain. RidL does not seem to account for the entire effect of mitochondrial fragmentation, since the mitochondria in Δ*icmT*-infected cells were still more elongated, as observed previously ^11^. Taken together, these results indicate that the *L. pneumophila* effector protein RidL promotes mitochondrial fragmentation.

**Fig. 5.**
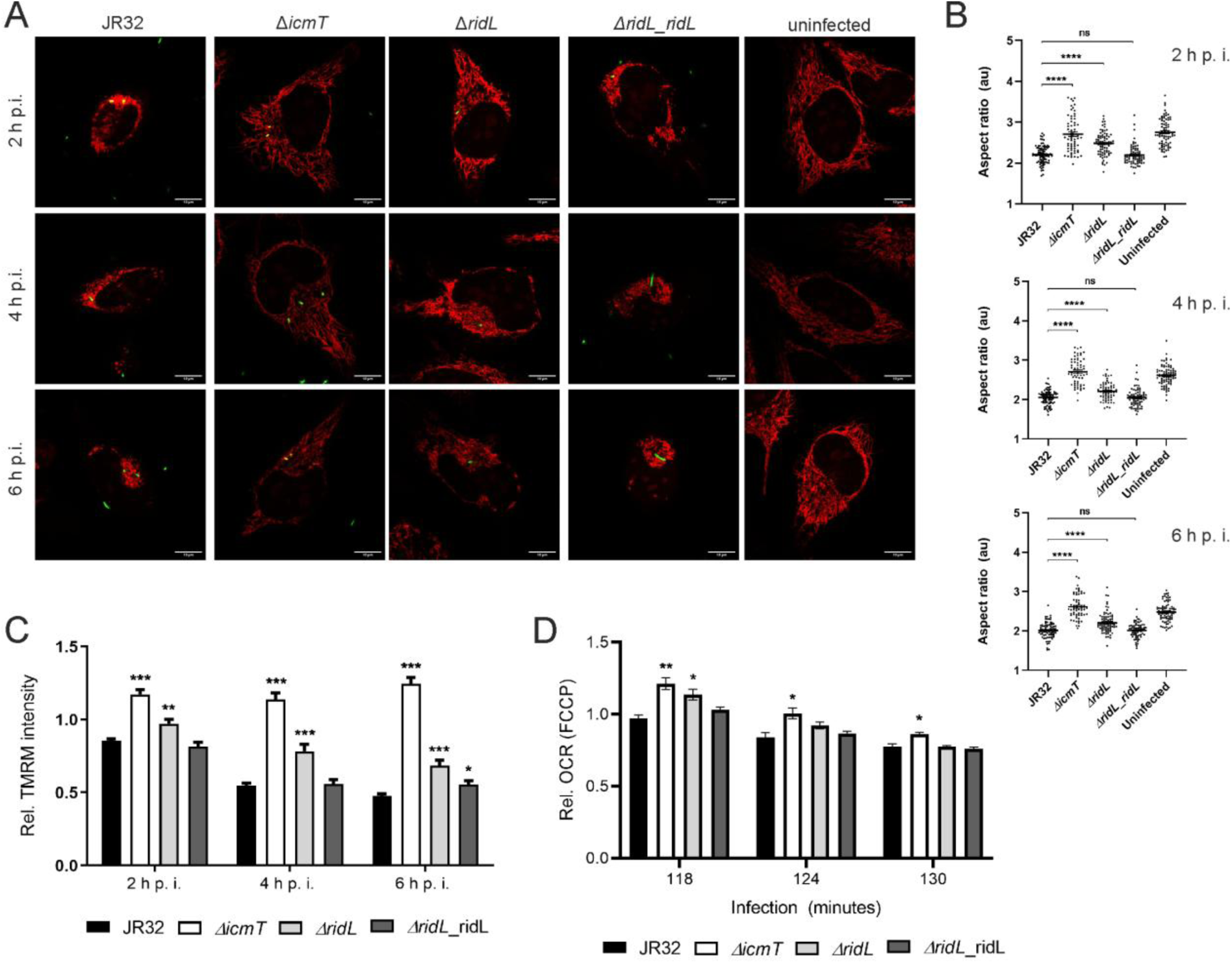
**During *L. pneumophila* infection RidL promotes fragmentation and impairs function of mitochondria**. (**A**) HeLa cells were stained with MitoTracker Orange CMTMRos (100 nM, 30 min, 37°C) and infected (MOI 100, 2-6 h) with GFP-producing *L. pneumophila* JR32, *ΔicmT*, *ΔridL*, or *ΔridL*_*ridL* (pNT28 or pIF009). Mitochondrial morphology was assessed by confocal laser scanning microscopy at the time points post infection (p.i.) indicated. Scale bars, 10 μm. Images are representative of 3 independent experiments. (**B**) Quantification of (A). In infected cells, the mitochondrial aspect ratio (d_max_ / d_min)_ was assessed using the Mitochondria Analyzer plugin for Image J. Dots represent all analyzed cells from three independent experiments (20-30 cells each; Student’s *t*-test, ****, p < 0.0001). (**C**) HeLa cells were infected (MOI 100, 2-6 h) with GFP-producing *L. pneumophila* JR32, *ΔicmT*, *ΔridL*, or *ΔridL*_*ridL* (pNT28 or pIF009) and stained with the membrane potential probe tetramethylrhodamine methyl ester (TMRM; 20 nM, 30 min, 37°C). GFP and TMRM fluorescence were assessed by flow cytometry at the time points p.i. indicated. Graphs show means + SEM of TMRM intensities of infected cells relative to uninfected cells of nine samples each from three independent experiments (> 15.000 cells per sample; Student’s *t*-test, *, p < 0.05; **, p < 0.01; ***, p < 0.001, relative to strain JR32). (**D**) HeLa cells were infected (MOI 10) with *L. pneumophila* strains JR32, Δ*icmT*, Δ*ridL* or Δ*ridL_ridL* (pNT28 or pIF009), subjected to a mitochondrial stress test, and the oxygen consumption rate (OCR) was measured by a Seahorse XF Pro analyzer. Oligomycin (final concentration 1 μM), FCCP (0.5 μM) and rotenone / antimycin A (0.5 μM) were injected at the indicated time points (see **Fig. S6c**). Graphs show relative OCR values upon injection of oligomycin and FCCP (normalized to OCR before mitochondrial stress test). Data taken from **Fig. S6c**; means + SEM of three independent experiments (Student’s *t*-test, *, p < 0.05, **, p < 0.01).

To assess the functional implications of RidL-dependent mitochondrial fragmentation, we tested the mitochondrial membrane potential and mitochondrial respiration (oxygen consumption rate) of *L. pneumophila*-infected HeLa cells. Along with mitochondrial fragmentation, RidL was found to also promote the dissipation of the mitochondrial membrane potential (**Figs. 5C**, **S6B**). Compared to the wild-type strain, the Δ*ridL* mutant strain was significantly less effective in reducing the mitochondrial membrane potential of infected cells, and the complemented strain reached wild-type levels of membrane potential dissipation.

Mitochondrial respiration was impaired upon infection of HeLa cells with *L. pneumophila* wild-type, but less so with the Δ*ridL* mutant strain (**Figs. 5D, S6C**). This phenotype was particularly pronounced upon addition of the uncoupler carbonyl cyanide-p-trifluoromethoxyphenyl-hydrazone (FCCP) to assess maximal respiratory capacity, and the complemented Δ*ridL* mutant strain reached wild-type levels of respiration inhibition. In contrast, upon infection of HeLa cells with the *L. pneumophila* Δ*icmT* mutant strain, the mitochondria were not fragmented and neither the mitochondrial membrane potential nor respiration were affected. In summary, these results indicate that in *L*. *pneumophila*-infected cells, RidL contributes to mitochondrial fragmentation, impairs the mitochondrial membrane potential and respiration.

### RidL promotes phosphorylation activation of Drp1 in *L. pneumophila*-infected cells

Drp1 activity is regulated in a complex manner by posttranslational modifications, including activation by phosphorylation at Ser616 ^19^. Since RidL promotes mitochondrial fission and impairs mitochondrial functions, we hypothesized that RidL might activate Drp1, perhaps by promoting phosphorylation at Ser616. To test this hypothesis, we infected HeLa cells with mCerulean-producing *L. pneumophila* strain JR32, Δ*icmT*, Δ*ridL*, complemented Δ*ridL* mutant, or Δ*ridL* mutant producing RidL_Δβ-loop_ lacking the retromer-interacting β-loop and performed confocal microscopy using a phospho-Ser616 specific anti-Drp1 antibody (**Figs. 6A, S7**). This approach revealed that upon infection with *L. pneumophila* wild-type, approximately 50% of the detected Drp1 localized with the mitochondrial stain MitoTracker, while this was the case for less than 10% or 20% of the detected Drp1 upon infection with *L. pneumophila* Δ*icmT* or Δ*ridL*, respectively (**Fig. 6B**). Co-localization of Drp1 with MitoTracker was restored by supplying plasmid-produced RidL or RidL_Δβ-loop_.

**Fig. 6.**
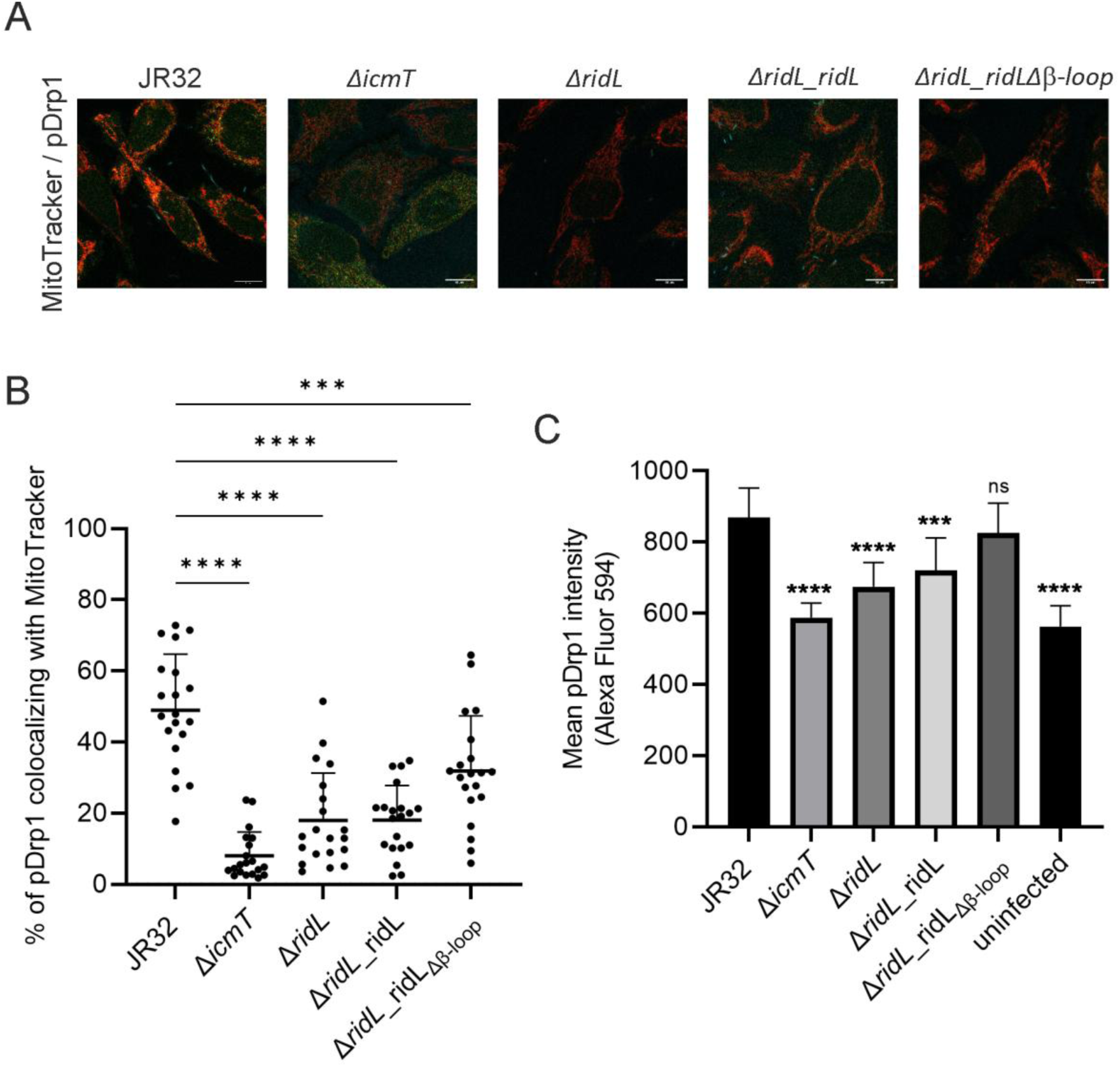
RidL promotes phosphorylation activation of Drp1 in *L. pneumophila*-infected cells. (**A**) HeLa cells were treated with MitoTracker Deep Red (30 min), infected (MOI 50, 2 h) with mCerulean-producing *L. pneumophila* strains JR32, Δ*icmT*, Δ*ridL*, Δ*ridL_ridL*, or Δ*ridL_ridL*Δ*β-loop* (pNP99, pKB208, or pKB209), immuno-stained with an anti-phospho-Drp1 antibody (Ser616) followed by an Alexa Fluor 488-coupled secondary antibody and analyzed by fluorescence microscopy. (**B**) Quantification of co-localization of MitoTracker and Drp1 (A) is shown (means and SD). (**C**) HeLa cells were treated with MitoTracker Deep Red (30 min), infected (MOI 50, 2 h) with GFP-producing *L. pneumophila* strains JR32, Δ*icmT*, Δ*ridL*, Δ*ridL_ridL*, or Δ*ridL_ridL*Δ*β-loop* (pNT28, pIF009, or pKB209), immuno-stained with an anti-phospho-Drp1 antibody (Ser616) followed by an Alexa Fluor 488-coupled secondary antibody and analyzed by flow cytometry. Means and standard deviation of three independent experiments are shown, all samples compared to JR32 by one-way ANOVA (***, p < 0.001; ****, p < 0.0001).

To validate these results, we performed an analogous experiment using flow cytometry (**Fig. 6C**). To this end, HeLa cells were infected with GFP-producing *L. pneumophila* strain JR32, Δ*icmT*, Δ*ridL*, complemented Δ*ridL* mutant, or Δ*ridL* mutant producing RidL_Δβ-loop_ lacking the retromer-interacting β-loop, fixed and analyzed by flow cytometry. This approach indicated that compared to cells infected with *L. pneumophila* wild-type the mean fluorescence intensity of cells infected with *L. pneumophila* Δ*icmT* or Δ*ridL* was significantly lower (**Fig. 6C**). Again, the phenotype was complemented by supplying plasmid-produced RidL or RidL_Δβ-loop_. Taken together, these results revealed that RidL promotes the activating phosphorylation of Drp1 at the amino acid residue Ser616. Intriguingly, this activity was observed for RidL as well as for RidL_Δβ-loop_, indicating that binding to the retromer is not required for Drp1 activation.

## Discussion

In the present study we show that the *L. pneumophila* effector protein RidL binds through a C-terminal fragment to the large dynamin-like mitochondrial fission GTPase Drp1 (**Fig. 1**). *In vitro*, RidL impairs GTPase activity (**Fig. 1**) and inhibits Drp1 oligomerization and spiralization (**Fig. 2**). Upon ectopic production in yeast, RidL interacts with several closely related and evolutionary conserved large fission GTPases (**Fig. 3**). Finally, in *L. pneumophila*-infected host cells, RidL associates with mitochondria and increases the levels of mitochondrial Drp1 as well as Tom20 (**Fig. 4**), promotes mitochondrial fragmentation and impairs mitochondrial function (**Fig. 5**), and upregulates phosphorylation of Drp1 (**Fig. 6**). While the N-terminal fragment of RidL has been characterized to interact with the Vps29 subunit of the cargo recognition subunit of the retromer coat complex, thereby displacing the Rab7 GAP TBC1D5 ^30–33^, the interactor(s) and function of the C-terminal fragment of RidL were unknown. Here we reveal that the C-terminal fragment of RidL binds to Drp1 *in vitro* (**Fig. 1F**) and localizes to mitochondria upon ectopic production (**Fig. 4CD**).

Drp1 is the major fission GTPase promoting mitochondrial fragmentation ^14^. Upon secretion of RidL in *L. pneumophila*-infected cells, significantly more Drp1 (**Fig. 4G**) or Tom20 (**Fig. 4H**) localized to mitochondria. The mechanism underlying the RidL-dependent mitochondrial fragmentation likely involves the activation of Drp1 by Ser616 phosphorylation (**Fig. 6**). Activation phosphorylation might either occur in the cytoplasm followed by recruitment of activated Drp1 to mitochondria, or by phosphorylation of mitochondria-bound Drp1. Several eukaryotic kinases modulate the activity of Drp1 ^14,19^, but the kinase activating Drp1 in the context of *L. pneumophila* infection has not been identified. Moreover, neither bioinformatics nor experimental evidence was obtained that RidL itself in a kinase. Our results are in agreement with the notion that RidL might either recruit Drp1 to mitochondria and/or prevent the removal/turnover of Drp1 from mitochondria.

Alternatively, or additionally to the direct mitochondrial recruitment or stabilization of Drp1 or phospho-Drp1, the mechanism underlying RidL-dependent mitochondrial fragmentation might involve the modulation of Drp1 removal/turnover via MDV formation. Mitochondrial protein turnover occurs – at least partly – hrough the formation of MDVs ^25^. While the mitochondrial recruitment of Drp1, oligomerization, and membrane constriction do not involve the retromer coat complex, Drp1 is indeed an MDV cargo in a pathway implicating the retromer ^26,27^. Furthermore, Drp1 is also recruited to the sites of nascent MDVs and promotes MDV fission ^24^. RidL inhibits the retromer in endosome-derived retrograde trafficking ^30,31^, and analogously, RidL might inhibit the (initial step of) MDV formation and removal of (inactive) Drp1 via this route. However, RidL promotes the accumulation of Drp1 and phospho-Drp1 on mitochondria (**Figs. 5, 6**), and therefore, the (final steps of) MDV formation, i.e. the fission of MDVs from mitochondria, might be eased. Future studies will shed light on whether and how MDV formation is implicated in RidL-mediated mitochondrial fragmentation.

Our study provides novel links between the retromer and Drp1, mitochondrial dynamics and MDV trafficking. In *L. pneumophila*-infected cells, RidL promotes the mitochondrial accumulation of not only Drp1 (**Fig. 4G**) but also Tom20 (**Fig. 4H**). Accordingly, RidL might interfere with the general protein turnover and quality control of mitochondria. Further studies will explore the role of RidL for MDV trafficking in infection biology and will also use RidL as a tool to dissect the role of MDV trafficking in cell biology, neurodegeneration and cancer.

Intriguingly, RidL is not the only *L. pneumophila* effector protein targeting the MDV pathway. The Ran GTPase activator LegG1/MitF stabilizes microtubules ^10,12^, likely accounting for its effect on promoting mitochondrial fragmentation ^11^ and the microtubule-dependent formation of mitochondria-derived tubules ^25^. Moreover, the effector Lpg1137 cleaves the SNARE syntaxin 17 ^9^, thereby likely inhibiting the final step of MDV fusion with lysosomes^28^. Several other *L. pneumophila* effectors have been shown to target mitochondria, such as the nucleotide carrier protein LncP ^6^, or the ADP/ATP translocase ADP-ribosyltransferase Lpg0080/Ceg3 and the antagonist hydrolase Lpg0081 ^7,8^. In summary, *L. pneumophila* seems to adopt a “double-hit” strategy to impair mitochondrial dynamics and function. The pathogen not only directly targets mitochondrial components but might also interfere with their turnover. RidL lacking the Vps29-binding β-loop but not wild-type RidL bound Drp1 in lysates of HeLa cells (**Fig. 1BC**), and full-length RidL bound Drp1 more strongly than the C-terminal fragment RidL_C_ (RidL_437-1100_) (**Fig. 1F**). Accordingly, the Vps29-binding β-loop and the N-terminal fragment of RidL seem to play a role for the interaction of the effector with the large GTPase. Perhaps full-length RidL interacts with oligomeric Drp1, while the C-terminal of RidL fragment does not. Moreover, it is unclear whether and how the interaction of RidL with Vps29 and/or the entire retromer cargo recognition complex affects its binding to Drp1. In agreement with a mutual dependency of different target protein interactions, the predicted structure of RidL comprises the Vps29-binding “foot” and “leg” domains, linked by an apparently flexible hinge to the Drp1-binding “arm” and “fist” domains (**Fig. 1A**). Intriguingly, RidL_Δβ-loop_ promotes the production of phospho-Drp1 on mitochondria, and therefore, binding to the retromer seems dispensable for this activity (**Fig. 6**). Finally, the presence of a lipid bilayer membrane likely also plays an important role for the Vps29-RidL-Drp1 interactions. Of note, RidL binds to the phosphoinositide lipid PtdIns(3)*P* ^30^, which might also affect the interactions among the effector, Vps29 and Drp1. Further mechanistic studies will address in detail these protein-protein and protein-lipid interactions.

*L. pneumophila* invests considerable resources to subvert mitochondria. An efficient inhibition of mitochondrial function likely provides an advantage for the strictly aerobic pathogen *L. pneumophila* regarding the competition between mitochondria and the bacteria for intracellularly available nutrients. In agreement with this notion, cellular lipid droplets associate with LCVs and are used by *L. pneumophila* as a source of catabolic fatty acids ^44^. Further studies will address the intricate (competitive) relationship between *L. pneumophila* and mitochondria.

## Materials and Methods

### Cells, bacteria, and infection

Human HeLa and A549 lung epithelial carcinoma cells were cultivated in RPMI 1640 medium, supplemented with 10% fetal bovine serum (FBS) and 2 mM L-glutamine (all from Gibco; ThermoFisher Scientific). HEK293 cells were cultivated in Dulbecco’s Modified Eagle Medium (DMEM; Gibco), supplemented with 10% FBS and 2 mM L-glutamine. The cells were maintained at 37°C and 5% CO_2_ in a humid atmosphere.

*L. pneumophila* strains harboring derivatives of pMMB207-C (pIF009, pKB198, pKB208, pNP99, pNT28) were grown on charcoal yeast extract (CYE) agar plates, buffered with *N*-(2-acetamido)-2-aminoethane sulfonic acid (ACES) containing 5-10 μg/ml chloramphenicol (Cam) at 37°C for 3 days. Bacterial over-night cultures were prepared by resuspending the strains in AYE containing 5 μg/ml Cam (OD_600_ of 0.1) and growing the liquid cultures at 37°C for 20.5-21.5 h until an OD_600_ of ca. 5 was reached.

For infection of mammalian cells, bacterial suspensions were prepared in the appropriate mammalian cell growth medium and added to the cells at the multiplicity of infection (MOI) indicated. The infections were synchronized by centrifugation (450 *g*, 10 min). The infected cells were washed 1 h post infection with DPBS (Gibco) and further incubated in fresh growth medium for the time indicated.

### Molecular cloning

Plasmids and oligonucleotides used in this study are listed in **Table S1** and **Table S2**, respectively. If not indicated otherwise, PCRs were performed using Phusion High-Fidelity DNA polymerase (Thermo Scientific) according to the manufacturer’s protocol and with addition of DMSO. Backbones and inserts were digested with restriction enzymes from ThermoFisher Scientific or New England Biolabs (NEB) at 37°C for 20-60 minutes. Ligations were performed at a vector:insert ratio of 1:3 with T4 DNA Ligase (NEB) at either 4°C over-night or at room temperature (RT) for 1 h. Competent *E. coli* TOP10 were transformed with the ligated constructs through heat shock (42°C, 30 sec.). Transformants were selected on LB plates containing ampicillin (Amp) or kanamycin (Kan). Colony PCRs were performed with inhouse Phusion or Taq polymerase. All PCR-amplified constructs were sequenced (Microsynth). Plasmids were isolated using the NucleoSpin Plasmid Mini kit for plasmid DNA (Macherey-Nagel) or with the EndoFree Plasmid Maxi Kit (Qiagen). PCR products were extracted from gels with the NucleoSpin Gel and PCR Clean-up Kit (Macherey-Nagel).

For the generation of yeast two-hybrid (Y2H) vectors, RidL was amplified from template pKB002 using the primers oKA001/oLS163 and fused to the DNA-binding domain of pLexA (BamHI/SalI), resulting in pKA001. Large GTPases were fused to the activating domain of pAct2.2. Vps1 was amplified using oKA004/oKA005 from template pKB242 and inserted into pAct2.2 through NdeI/EcoRI, resulting in pKA004. Drp1 was amplified with oKA020/oKA021 from template mCh-Drp1 (Addgene #49152 ^45^) and inserted via NdeI/BamHI, yielding pKA009. Dnm2 was amplified with oKA023/oKA024 from template pKB241 and inserted into pAct2.2 through EcoRI/NdeI, resulting in pKA011. Dnm3 was amplified with oKA261/oKA262 from template pDONR201-DNM3 (HsCD00080526; DNASU Plasmid Repository) and inserted into pAct2.2 via SmaI/BamHI, yielding pKA104. The construct pAct2.2-Dnm1 (pKA103) was assembled through NEBuilder HiFi DNA Assembly (NEB). The insert (*Dnm1*) was amplified from pEGFP-N1-dynamin (Addgene #120313 ^46^) using primers oKA288/oKA289 and Q5 Hot Start High-Fidelity DNA Polymerase (NEB) including Q5 High GC Enhancer. Due to the lack of matching restriction sites, backbone pAct2.2 was fully amplified using oKA290/oKA291 and Q5 Hot Start High-Fidelity DNA Polymerase and DpnI-digested for clearance of the DNA template. DNA assembly was performed as indicated in the NEBuilder HiFi DNA Assembly manufacturer’s manual. Both, the DNA-binding domain of pAct2.2 and the insert *Dnm1* were sequenced. For pACT2.2-DymA (pKA014) *DymA* was amplified with oKA029/oKA030 from pDXA-GFP-DymA (gift from F. Letourneur), backbone and insert were digested with NdeI/EcoRI, and the backbone was treated with CIAP (Calf Intestinal Alkaline Phosphatase; Invitrogen) before ligation. For pAct2.2-DymB (pKA015) *DymB* was initially amplified with oKA263/oKA264 from pME18SFL3-DymB (#G01672; National BioResource Project (NBRP), Nenkin, Japan). The PCR product was cleaned up and conducted to A-tailing (0.2 mM ATP, NEB One Taq Hot Start Polymerase, 94°C for 30 sec, 65°C for 10 min), followed by DpnI digest (37°C for 1 h, heat-inactivation at 80°C for 20 min) and ligated with the pGEM-T Easy vector (pGEM-T Easy Vector Systems; Promega) over-night at 4°C, resulting in the shuttle vector pKA105. DymB was cut out from pKA105 with NdeI/BamHI and ligated with pAct2.2, yielding pKA015.

To generate split-GFP vectors for bimolecular fluorescence complementation (BiFC) of RidL and large fission GTPases, RidL was amplified with the primers oKA183/oKA051 from template pKA001 and fused to GFP_11_-mCherry in pRS315-NOP1pr-GFP11-mCherry-PUS1 (pSJ1321, Addgene #86413 ^47^) by replacing PUS1 (NheI/SalI), resulting in pKA067. The large fission GTPases Drp1, Dnm1, Dnm2, Dnm3, DymA, DymB and Vps1 were fused to GFP_1-10_ in pRS316-NOP1pr-GFP1-10-SCS2TM (pSJ2039, Addgene #86418 ^47^) by replacing SCS2TM. For cloning of pKA100 (pRS316-GFP_1-10_-Drp1), pKA097 (pRS316-GFP_1-10_-Dnm1), pKA098 (pRS316-GFP_1-10_-Dnm2), pKA099 (pRS316-GFP_1-10_-Dnm3), pKA114 (pRS316-GFP_1-10_-DymA) and pKA072 (pRS316-GFP_1-10_-Vps1) the backbone was amplified (excluding the *scs2tm* sequence) with the primers oKA207/oKA208 and Q5 Hot Start High-Fidelity DNA Polymerase. *Drp1* was amplified with oKA245/oKA246, *Dnm1* with oKA239/oKA240, *Dnm2* with oKA241/oKA242, *Dnm3* with oKA243/oKA244, *DymA* with oKA280/oKA281 and *Vps1* with oKA209/oKA210. Inserts and fragments were assembled with NEBuilder HiFi DNA

Assembly (NEB). For generation of pRS315-GFP_1-10_-DymB (pKA115), an AgeI restriction site was initially introduced into the backbone pSJ2039, replacing *scs2tm*, by amplifying the full backbone excluding *scs2tm* with oKA224/oKA225. The PCR product was annealed with an oligonucleotide bridge containing the AgeI restriction site (oKA300) through NEBuilder assembly, resulting in plasmid pKA118. An AgeI-flanked *DymB* construct was amplified with oKA298/oKA299 and inserted into an pGEM-T Easy vector as described for pKA105, resulting in pKA117. Finally, DymB was cut out of pKA117 and inserted into pKA118 (AgeI), resulting in pKA115.

pKB248, leading to the production of GFP-RidL^opt^ was cloned by amplification of codon-optimized RidL from template pKB245 (pEGFP-N1-ridL^opt^; Genscript) with oKB160/oKB161 and insertion into pEGFP-C1 through Sal/BamHI. For construction of pKB249 (pEGFP-C1-ridL^opt^ -Stop) or pKB250 (pEGFP-C1-ridL^opt^_259-1167_), RidL fragments were amplified using pKB245 as the template and oKB162/oKB163 (pKB249) or oKB153/oKB160 (pKB250), respectively, and inserted into pEGFP-C1 through SalI/BamHI.

For production and purification of RidL fragments, RidL_1-258_ (oKB001/oKB082) and RidL_437-1100_ (oKB210/oKB179) fragments were amplified using pKB002 as a template and inserted into pINIT through an FX cloning system. RidL_1-258_ was cut (BspQI) and inserted into pBXNH3, resulting in pKB139, RidL_437-1100_ was inserted into pBXC3GH, resulting in pKB306. For Drp1 production, human Drp1 was amplified with oKA355/oKA356 using mCh-Drp1 or pKA080 as templates and inserted into pET21-Drp1 (Addgene #72927 ^48^) through NdeI/BamHI, replacing the murine Drp1, and yielding pKA173 (pET21-Drp1). The Drp1_A395D_ point mutation was created using QuikChange, with pKA173 as a template, yielding pEV001.

### Protein production and purification

To produce human Drp1, *E. coli* BL21(DE3) harboring plasmid pKA173 was grown in LB broth containing 100 μg/ml Amp, induced with 1 mM isopropyl-β-d-thiogalactoside (IPTG; Carl Roth) at OD_600_ 0.8 and left to grow overnight at 21°C. To produce RidL and its N- and C-terminal fragments, *E. coli* MC1061 containing a pBXNH3-derivative (N-terminal cleavable His_10_-tag) were grown in LB/100 μg/ml Amp and induced at OD_600_ 0.7 with 0.02% (w/v) L(+)-arabinose overnight at 16°C.

Bacterial pellets were resuspended on ice in lysis buffer (50 mM Tris/HCl, pH 7.5, 200 mM NaCl) supplemented with 3 mM MgSO_4_, 1 mM PMSF, and DNAse (Sigma) and lysed by using high-pressure homogenization (Microfluidizer; Microfluidics) with a pressure of 21,000 PSI. The resulting lysate was centrifuged (16’000 *g*, 4°C, 30 min). Lysates were then passed through a Ni^2+^-NTA affinity chromatography column, washed with wash buffer (50 mM imidazole, pH 7.5, 200 mM NaCl, 10% glycerol), and His-tagged protein was eluted with elution buffer (washing buffer with 300 mM imidazole). To remove imidazole, the buffer was exchanged with the size-exclusion chromatography (SEC) buffer (50 mM Tris/HCl, pH 7.5, 200 mM NaCl) using PD10 desalting columns (GE Healthcare). The protein was incubated overnight with 3 C protease to remove the His-tag (produced in-house, 10 ug/ml). As a final purification step, the protein was loaded onto a Superdex 200 10/300 GL (GE Healthcare) equilibrated with SEC buffer. Fractions where the monodispersed peak eluted were combined and concentrated with Amicon Ultra-4 centrifugal filter units, using a molecular weight cutoff of 10 (RidL-N), 50 (RidL-C, Drp1) or 100 kDa (full-length RidL). Protein concentration was determined by OD_280_ using a NanoDrop 2000 photospectrometer and calculated based on theoretical extinction coefficients (www.expasy.ch/tools/protparam.html). For storage, the protein aliquots were snap-frozen and stored at −80°C.

### GTPase activity assay

A continuous GTPase activity assay was used to analyze the activity of Drp1 in the presence and absence of RidL. Prior to running the assay, Drp1 samples with and without RidL were incubated for 1 h at RT, with shaking at 350 rpm. Drp1 was diluted to a final concentration of 1 μM in a master mix solution (50 mM HEPES/KOH, pH 8, 150 mM NaCl, 0.5 mM MgCl_2_, 10 mM β-mercaptoethanol, 4 mM phosphoenolpyruvate, 0.3 mM NADH, and 10 units of pyruvate kinase and lactate dehydrogenase). Once the samples were pipetted into a 96-well UV-transparent plate, GTP was added to all samples to a final concentration of 1 mM. The depletion of NADH was measured at 340 nm (BioTek Cytation 5; Agilent) over the course of 2 h. As controls, wells were filled with master mix with and without NADH and used to correct for background. The rate of NADH depletion was used to measure GTPase activity of Drp1.

### Negative stain electron microscopy

Drp1 was visualized by negative stain electron microscopy as described ^49^. To induce filaments, Drp1 was diluted into negative staining buffer (50 mM Tris/HCl, pH 7.5, 200 mM NaCl, 1 mM DTT) and incubated with 0.5 mM GMPPCP for one hour. Samples incubated with RidL and SidC were added simultaneously upon the addition of GMPPCP. To visualize the filaments, samples were placed onto carbon film coated, 300 mesh, copper grids (Science Services). After 1 min of incubation, the sample was blotted away with filter paper. The grids were immediately washed with uranyl acetate, blotted again, and then incubated with the uranyl acetate once again for 1 min After staining, the grids were dried for about 10 min before imaging on the FEI Tecnai Spirit, a 120 kV electron microscope from the University of Zürich Center for Microscopy (ZMB).

### Dynamic light scattering

To quantify the presence and absence of Drp1 filaments in solution, we measured the amount of scattered light among various samples using dynamic light scattering (Zetasizer Nano ZS; Malvern Panalytical). The scattered intensity was measured at 173 degrees. The oligomerization of Drp1 (1 µM) induced by 0.5 mM GMPPCP, in the absence or presence of RidL (3 µM) or SidC (3 µM), was assessed in a volume of 100 µl. Immediately after the components were mixed, the sample was pipetted into a cuvette, and scattered intensity was read by the machine in 1-min intervals. The hydrodynamic size distributions were calculated using cumulant and CONTIN methods.

### Yeast two-hybrid assays

For yeast two hybrid (Y2H) assays, RidL was fused to a LexA DNA-binding domain (pLexA), and large fission GTPases (Drp1, Dnm1, Dnm2, Dnm3, DymA, DymB or Vps1) were fused to the Gal4 activation domain (pAct2.2). Upon transformation of the Y2H reporter strain NMY32, an interaction between bait and prey proteins leads to the reversion of histidine auxotrophy.

Strain NMY32 was grown on yeast-extract peptone dextrose (YPD) agar plates at 30°C for 3 days. For the generation of competent cells, NMY32 was grown in liquid YPD at 30°C over-night, diluted to an OD_600_ of 0.2 and further incubated until OD_600_ of 0.6-0.8 was reached. Yeast cells were harvested (600 *g*, 3 min, 4°C), washed with bi-distilled H_2_O (ddH_2_O) and LiSorb (10 mM Trizma base, pH 8.0, 100 mM C_2_H_3_LiO_2_, 1 mM EDTA, 1 M sorbitol) and finally resuspended in LiSorb.

For transformation, 500 ng plasmid DNA, 50 μg sheared salmon sperm DNA (Invitrogen), 50 μl of competent yeast cells and 300 μl LiPEG (10 mM Trizma base, pH 8.0, 100 mM C_2_H_3_LiO_2_, 1 mM EDTA, 45% PEG4000) were incubated at 30°C for 30 min while being mixed. Then, 35 μl DMSO were added to the mixture before exposure to a heat-shock at 42°C for 15 min. Yeast transformants were washed with and resuspended in ddH_2_O, plated for selection onto SD-Leu-Trp agar plates and incubated for 3 days at 30°C.

For Y2H assays, single yeast colonies were re-streaked onto SD-Leu-Trp agar plates, grown for another 2-3 days at 30°C and resuspended in ddH_2_O. Yeast suspensions were adjusted to OD_600_ of 0.5 and spotted in a 10-fold dilution series onto SD-His agar plates containing 0.5 mM 3-amino-1,2,4-triazole (3-AT) and grown for 3 days at 30°C to select for protein-protein interactions. As a control, the serial dilutions were additionally spotted onto SD-Leu-Trp agar plates.

### Bimolecular fluorescence complementation

For bimolecular fluorescence complementation (BiFC) using split GFP, RidL was fused to one strand of the GFP β-barrel (GFP_11_) and mCherry using the plasmid pRS315-NOP1pr-GFP_11_-mCherry (derived from pSJ1321) ^47^, resulting in the fusion protein GFP_11_-mCherry-RidL (pKA067). Large GTPases (Drp1, Dnm1, Dnm2, Dnm3, DymA, DymB or Vps1) were fused to the GFP_1-10_ fragment by modification of pSJ2039 ^47^. The yeast strain BY4741 was first transformed with pKA067 (GFP_11_-mCherry-RidL), selected on SD-Leu, and competent BY4741/pKA067 was then transformed with plasmids encoding GFP_1-10_ fused to a large GTPase (selection on SD-Leu-Ura).

For fluorescence imaging of GFP- and mCherry signals, over-night cultures were prepared by inoculating selective media with the transformants grown at 30°C. On the day of the experiment, over-night cultures were diluted to an OD_600_ of 0.2 and further grown until an OD_600_ of 0.6-0.8 was reached. At this stage, aliquots of the yeast cultures were transferred to 8-well μ-slides, ibiTreat (#80826; ibidi), embedded with 0.1% agarose and imaged by confocal laser scanning microscopy (Leica SP8 DMi8 CS with AFC, objective HC PL APO CS2 63× oil) with 4× zoom, bi-directional laser scan and a scanning speed of 400 Hz. For each sample, 25 tile scans were recorded.

### Protein pulldown in cell lysates

For pulldown experiments, purified RidL (full-length, N-, and C-terminal fragments, RidL-Δβ) or SidC were covalently linked to polyacrylamide beads (UltraLink Biosupport; Thermo Scientific) according to the manufacturer’s protocol. Efficiency of the coupling was determined by using a BCA protein determination assay to check equimolar amounts of RidL being linked to the beads. To produce lysates, HeLa cells were seeded at a density of 3 × 10^6^ cells in T75 flasks and were left to grow for 24 h as described above. The cells were collected and lysed with a ball homogenizer (Isobiotec) using SEM buffer (250 mM sucrose, 1 mM EDTA, 10 mM MOPS-KOH, pH 7.2) containing protease inhibitors (Pierce Protease Inhibitor Mini Tablets, EDTA-free; ThermoFisher Scientific). Lysed cells were collected and spun down at 16,000 *g* at 4°C to separate proteins from cell debris. The supernatant was collected and incubated with the RidL or SidC-coupled beads for 1 h at 4°C. After incubation, the liquid was removed using a syringe, the beads were washed, collected, resuspended in SDS buffer, and boiled at 95°C for 5 min, prior to analysis by SDS-PAGE.

### Western blot analysis

For assessment of RidL, Drp1, Vps29, AIF and GAPDH in cell lysates, isolated mitochondria and other cellular fractions, protein samples were separated by SDS-PAGE (10% acrylamide gels) and blotted onto nitrocellulose membranes at 100 V, 4°C for 1 h. In general, membranes were blocked in 5% milk or 5% BSA (Drp1) for 30-60 min at RT and incubated over-night at 4°C with primary antibodies: polyclonal anti-AIF (17984-1-AP, 1:5000; Proteintech), rabbit mAb anti-GAPDH (14C10) (#2118, 1:1000; Cell Signaling Technology), mouse anti-Vps29 (#sc-398874, 1:100; Santa Cruz Biotechnology), rabbit mAB anti-DRP1 (D8H5) (#5391, 1:1000; Cell Signaling Technology), mouse monoclonal anti-GFP Living Colors JL-8 (#632380, 1:5000; Clontech), or polyclonal anti-RidL (1:5000, ^30^). Secondary antibodies (#NA934-1ML, ECL donkey anti-rabbit IgG HRP-linked whole Ab; Cytiva/Amersham; #31430, goat anti-mouse IgG [H+L] HRP-linked secondary antibody; Invitrogen) were diluted 1:5000 in 4% milk and incubated for 1 h at RT. Protein detection was conducted with Westar Sun ECL Substrate (Cyanagen) and the ImageQuant 800 (Amersham).

### Mitochondria isolation

Mitochondria were isolated from infected HeLa or transfected HEK293 cells as described ^50^. In brief, previously infected or transfected cells from 12 confluent T75 flasks were trypsinized, collected (450 *g*, 5 min), resuspended in SEM buffer, supplemented with protease inhibitors (ThermoFisher Scientific) and lysed by using a ball homogenizer. This cell lysate was centrifuged (800 *g*, 4°C, 5 min), resulting in a nuclei-enriched pellet and an organelle-containing supernatant. The supernatant was centrifuged again (800 *g*, 4°C, 5 min), followed by a final centrifugation step (8’000 *g*, 10 min, 4°C), resulting in a mitochondria-enriched pellet (crude mitochondria, “cM”) and a supernatant referred as a cytoplasmic fraction (“cyt”). To receive a pure mitochondria fraction (“pM”), the crude mitochondria pellet was resuspended in SEM buffer containing protease inhibitor, subjected to sucrose density gradient ultracentrifugation (60%, 32%, 23% and 15% sucrose in 10 mM MOPS/1 mM EDTA, pH 8.0), and centrifuged (ca. 133’500 *g*, 1 h, 4°C). The pure mitochondria were retrieved from the interphase between 60% and 32% sucrose, diluted with the 2-fold volume of SEM buffer, collected (8’000 *g*, 10 min, 4°C) and finally resuspended in 200 μl SEM buffer.

For the expression of GFP-tagged full-length RidL and fragments, HEK293 cells were transfected in 90% confluent T75 flasks with each 23.8 μg plasmid DNA per flask by polyethylenimine treatment (PEI MAX; Polysciences). A mix of plasmid DNA, PEI (95 μg per flask, dissolved in H_2_O) and OptiMEM I (Gibco) was added to HEK293 cells for one day before being replaced by growth medium. Transfected HEK293 cells were further processed two days after transfection.

### Imaging of mitochondrial fragmentation

Mitochondrial networks were imaged and analyzed within individual infected HeLa cells. To this end, the cells were seeded at low densities (25’000 cells per well) into 24 well μ-plates (#82426; ibidi) and incubated for ca. 24 h. Before infection, cells were stained with 100 nM MitoTracker^TM^ Orange CMTMRos (ThermoFisher Scientific) for 30 min at 37°C, washed with DPBS and infected (MOI 100) with *L. pneumophila* strains JR32, *ΔicmT*, *ΔridL* harboring pNT28 (GFP) or *ΔridL* harboring pIF009 (GFP, RidL) as described above. 2-6 h post infection, the mitochondrial morphology was assessed by live confocal laser scanning microscopy (Leica SP8 DMi8 CS with AFC, objective HC PL APO CS2 63× oil) with 2× zoom, bi-directional laser scan and a scanning speed of 200 Hz. For quantification, mitochondria within individual cells were identified, and their mean branch lengths and aspect ratios (d_max_/d_min_, d = diameter) were assessed with the Mitochondria Analyzer plugin in Image J/Fiji ^51,52^.

### Membrane potential of infected cells

HeLa cells were seeded into 12-well plates the day before the experiment and grown at 37°C, 5% CO_2_ for ca. 24 h. Cells were infected (MOI 100) with *L. pneumophila* strains JR32, *ΔicmT*, *ΔridL* harboring pNT28 (GFP) or *ΔridL* harboring pIF009 (GFP, RidL) as described above. 1 h, 3 h and 5 h post infection, cells were detached by trypsinization and incubated with 20 nM tetramethyl rhodamine methyl ester (TMRM, ab275547; Abcam) diluted in growth medium (37°C, 5% CO_2_, 30 min). After incubation, cells were collected (450 *g*, 5 min), resuspended in DPBS and processed by flow cytometry with an LSR II Fortessa Cell Analyzer (BD Biosciences), corresponding to a total of 2 h, 4 h and 6h post infection. GFP-producing intracellular bacteria were assessed using the blue laser (488 nm), mirror 505LP and filter 530/30 at 450V, and the TMRM signal was assessed through the yellow/green laser (561) nm, mirror 600LP, filter 610/20 at 400V (FSC at 420V, SSC at 280V). > 10^4^ infected (GFP-positive) HeLa cells or the total sample volume was processed and measured. The mean TMRM intensities of all infected HeLa cells (based on the GFP-signal) were assessed with the FlowJo software. As controls, cells were treated with 50 μM carbonyl cyanide 3-chlorophenylhydrazone (CCCP, 98%; ThermoFisher Scientific or M20036, MitoProbe TMRM kit for flow cytometry; Invitrogen) for 5 min at 37°C, 5% CO_2_. TMRM was added to a final concentration of 20 nM directly to the CCCP-containing sample. The sample was further incubated (37°C, 5% CO_2_, 30 min) and processed as described above.

### Mitochondrial respiration and stress test

Mitochondrial basal and stressed respiration of *L. pneumophila*-infected HeLa cells were tested using a Seahorse XF Pro Analyzer (Agilent) and the Cell Mito Stress Test Kit (Agilent). To this end, 1.5 × 10^4^ HeLa cells per well were seeded into 96-well plates (Agilent), let rest for 1 h at RT, and incubated for one day at 37°C, 5% CO_2_. The basal respiration of uninfected HeLa cells was measured at 12 min and 6 min before infection (1 measurement cycle = 6 min). The HeLa cells were then infected (MOI 10, 450 *g*, 10 min) with *L. pneumophila* and subjected to respiration measurements for 15 cycles (6 min per cycle). The data was normalized to basal respiration rate. 100 min post infection, a mitochondrial stress test was performed according to the manufacturer’s protocol, with final drug concentrations of 1 μM oligomycin, 0.5 μM FCCP and 0.5 μM rotenone/antimycin A. The data was normalized to respiration rate of infected cells immediately before stress test.

### Protein depletion by RNA interference

The siRNA-mediated depletion of Dnm1 and Drp1 was performed as described ^53^. In brief, A549 cells were seeded and grown to ca. 80-90% confluency over-night before being transfected with siRNA oligonucleotides targeting Drp1 (Hs_DNM1L_4, Hs_DNM1L_8, Hs_DNM1L_9, Hs_DNM1L_10 FlexiTube siRNA; Quiagen), Dnm1 (FlexiTube GeneSolution GS1759 for DNM1; Qiagen) or Arf1 (Hs_ARF1_10 FlexiTube siRNA; Qiagen). The siRNA oligonucleotides were mixed with HiPerFect Transfection Reagent (Qiagen) and transferred into 96-well plates before adding each 2 × 10^4^ A549 cells. Transfected cells were incubated at 37°C, 5% CO_2_ for 48 h before infection with *L. pneumophila* JR32 or Δ*icmT* (MOI 10).

Bacterial GFP-signal was measured with a plate reader 1 h and 24 h post infection. To assess the cytotoxicity of protein depletion, the samples were stained with Zombie Aqua dye (1 h, 37°C) and processed by flow cytometry. The efficiency of protein depletion was checked by Western blot.

### Immunofluorescence microscopy

For immunostaining of Tom20, Drp1, and pDrp1 (Ser616), 10^5^ HeLa cells each were seeded on coverslips in 24-well plates overnight. The cells were stained with 100 nM MitoTracker Deep Red FM (ThermoFisher Scientific) for 30 min at 37°C, washed and infected (MOI 25, 2 h for Tom20 and Drp1, MOI 50 for 1h for pDrp1 (Ser616)) with mCerulean-producing *L. pneumophila* strains JR32, Δ*icmT*, or Δ*ridL* harboring pNP99 or Δ*ridL* harboring pKB208 or pKB209. Infected cells were fixed with 4% PFA (30 min, RT), permeabilized with 0.25% Triton X-100 (5 min, RT) and blocked with 10% FBS, diluted in DPBS or PBS (1 h, RT). Samples were incubated with either anti-Drp1 (C-terminal) polyclonal antibody (#12957-1-AP, 1:50 in 1% BSA; Proteintech), anti-Tom20 polyclonal antibody (#11802-1-AP, 1:200 in 1% BSA; Proteintech), or anti-phospho-Drp1 (Ser616) monoclonal antibody (#4494, 1:1000 in 1% BSA; Cell Signaling Technology) for 1.5 h at RT in the dark. After washing, samples were incubated with chicken anti-rabbit IgG (H+L) cross-adsorbed secondary antibody, Alexa Fluor 488 (#A-21441; ThermoFisher Scientific, 1:250 in 1% BSA) or F(ab’)2-goat anti-rabbit IgG (H+L) cross-adsorbed secondary antibody, Alexa Fluor 488 (A-11070; Invitrogen, 1:250 in 1% BSA) for 45 min. Coverslips were mounted with ProLong Diamond antifade mountant (ThermoFisher Scientific) and dried overnight at RT. Imaging was performed by confocal laser scanning microscopy (Leica SP8 DMi8 CS with AFC, objective HC PL APO CS2 63× oil) with 2× or 4× zoom, bi-directional laser scan and a scanning speed of 200 Hz. The samples were randomly picked without the GFP channel switched on. Unedited images were analyzed in Image J by defining individual thresholds for MitoTracker Deep Red, GFP and mCerulean channels and measuring mean signal intensities. For improved illustration, the brightness of fluorescence microscopy images was increased in Image J.

### Flow cytometry of immuno-stained cells

For flow cytometry of immuno-stained cells, 4 × 10^5^ HeLa cells were seeded per well of a 6-well plate and grown overnight. The cells were infected (MOI 50) with GFP-producing *L. pneumophila* strains JR32, Δ*icmT*, or Δ*ridL* haboring pNT28, or Δ*ridL* harboring pIF009 or pKB198. 1 h post infection, cells were washed two times with DPBS, detached by trypsinization for 3 min at 37°C, collected (500 *g*, 5 min) and fixed with 4% paraformaldehyde (15 min, RT). After one wash with PBS, cells were permeabilized with 90% methanol for 10 min on ice, followed by a wash with PBS. For immunostaining, cells were incubated (1 h, RT) with an anti-pDrp1 (Ser616) rabbit monoclonal antibody (#4494, 1:1000 in 0.5% BSA; Cell Signaling Technology), washed with PBS and incubated (30 min, RT) with an Alexa Fluor 594-conjugated secondary antibody (2 drops per mL of 0.5% BSA (according to the manufacturer’s protocol); R37117, Invitrogen). Controls were incubated in 0.5% BSA without the primary antibody. After a final wash step with PBS, cells were resuspended in PBS and processed by flow cytometry (BD LSRFortessa Cell Analyzer; BD Biosciences). Intracellular GFP-producing bacteria were assessed with the 488 nm laser (Blue) with mirror 505LP and filter 530/30 (channel Blue 530/30) at 380 V, pDrp1-bound Alexa Fluor 594-conjugated antibody signal was assessed with the 561 nm laser (Yellow/Green) with mirror 600LP and filter 610/20 (channel YG 610/20) at 470 V. Analysis was done with FlowJo software.

### Bioinformatics and statistics

Protein structure predictions were performed with the Alphafold2 algorithm ^54^. Each experiment was independently replicated at least three times, and representative images are shown. All statistical analyses were performed using GraphPad Prism version 7.01 for Windows, GraphPad Software, La Jolla California, USA (www.graphpad.com). Two-tailed Student’s *t* test was used, and significance levels are defined as follows: *, **, ***, or **** to indicate probability values of less than 0.05, 0.01, 0.001, or 0.0001, respectively.

## Acknowledgements

We would like to thank Daniela Portugal, Agnese Pisano, and Alexander Geiger for help with yeast work and the coupled GTPase assay. The plasmids pDXA-GFP-DymA and pME18SFL3-DymB were provided by François Letourneur and the National BioResource Project (NBRP) Nenkin, Japan, respectively. Electron microscopy and flow cytometry was performed at the University of Zürich (UZH) Center for Microscopy (ZMB) and the UZH Cytometry Facility. Research in the laboratory of H.H. was supported by the Swiss National Science Foundation (SNF; 175557, 207826). Research in the team of P.E. was supported by the French “Agence National de la Recherche” (ANR; grant ANR-21-CE15-0038-01) and the “Programmes Transversaux de Recherche” (PTR; grant PTR-651-23) from the Institut Pasteur. Research in the laboratory of C.B. was supported by the French Government (grant ANR-10-LABX-62-IBEID) and the “Fondation pour la Recherche Médicale” (grant EQU201903007847).

## Author contributions

Conceptualization: AK, ETV, VGP, HH

Methodology: AK, ETV, SH, KS, ALS, MR, FJG-R, TJ, XL

Investigation: AK, ETV, SH, KS, ALS, MR, FJG-R, TJ, XL

Visualization: AK, ETV, XL

Funding acquisition: CB, PE, VGP, HH

Project administration: HH Supervision: JAM, CB, PE, VGP, HH

Writing – original draft: AK, ETV, HH

Writing – review/editing: AK, ETV, SH, KS, ALS, MR, FJG-R, TJ, XL, JAM, CB, PE, VGP, HH

## Data and materials availability

All data are available in the main text or the Supplementary Information.

## Supplementary Information

Figures S1 to S7 Tables S1 and S2

Supplementary references

## Abbreviations

Drp1: dynamin-related protein-1
DLS: dynamic light scattering
EM: electron microscopy
Icm/Dot: intracellular multiplication/ defective organelle trafficking
LCV: *Legionella*-containing vacuole
MDV: mitochondria-derived vesicles
PI: phosphoinositide
T4SS: type IV secretion system.

## Supplementary Information

**Fig. S1.**
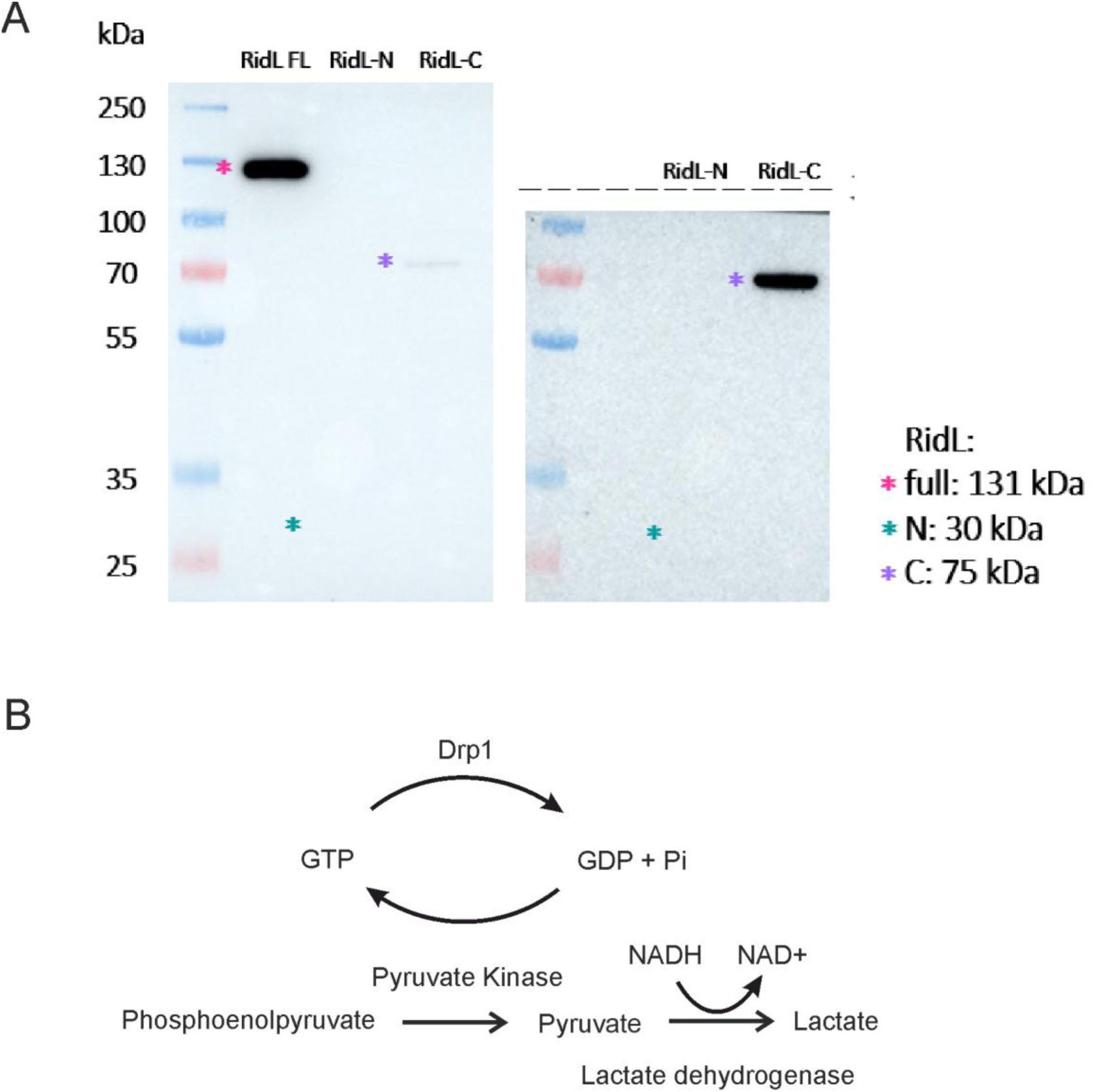
Coupled enzyme assay for Drp1 GTPase activity. (**A**) Purified RidL (131 kDa), RidL_N_ (RidL_1-258_, 30 kDa), or RidL_C_ (RidL_437-1100_, 75 kDa) were incubated with purified Drp1 (78 kDa) coupled to UltraLink Biosupport beads. The beads were washed once, and bound proteins were separated by SDS-PAGE and visualized by anti-RidL Western blot. Regular (left panel) and prolonged exposure (right panel) is shown. (**B**) Scheme of coupled enzyme assay for Drp1 GTPase activity, adapted from ^1^.

**Fig. S2.**
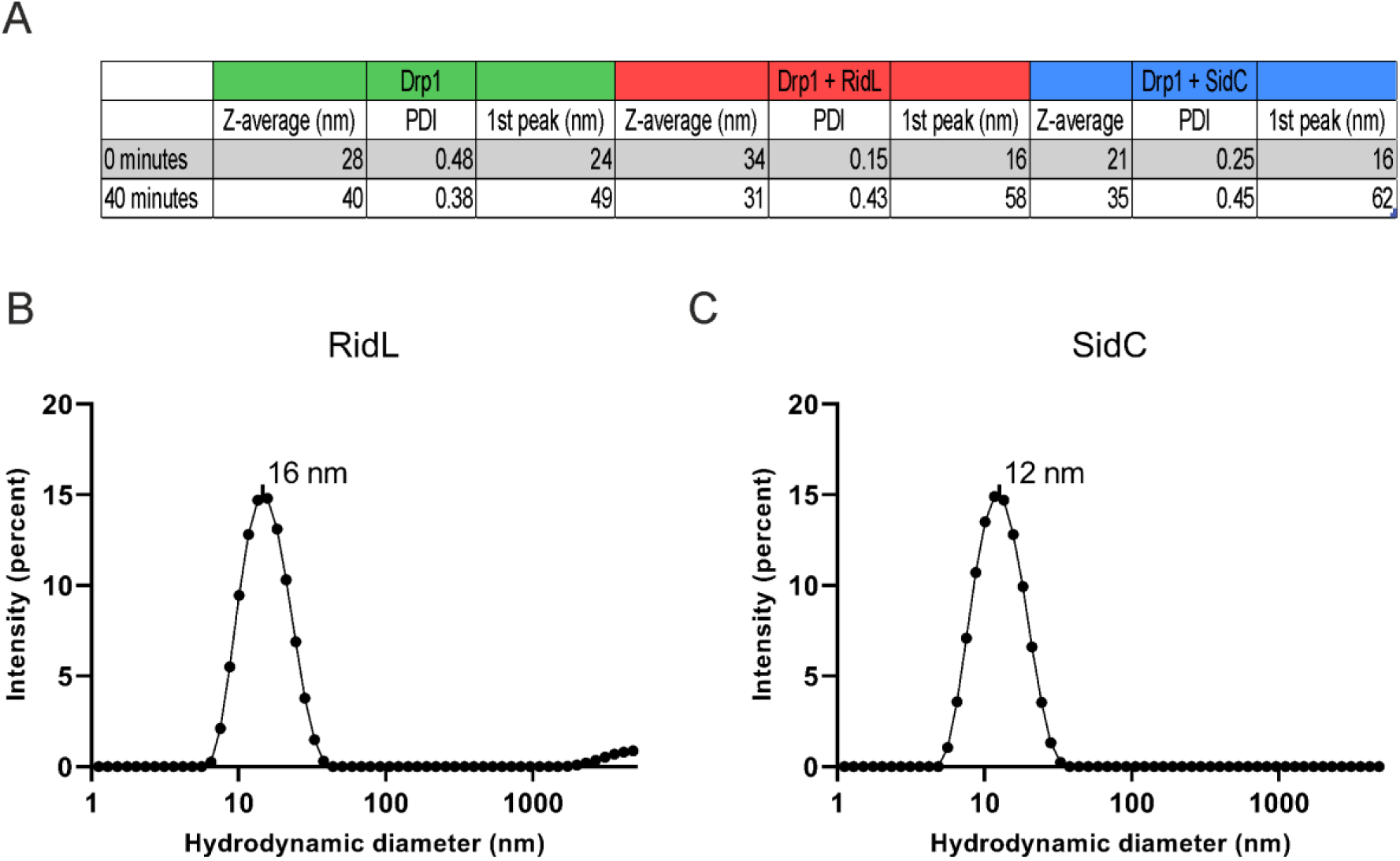
Dynamic light scattering values and profiles. (**A**) Hydrodynamic diameter (in terms of Z-average) and polydispersity index (PDI) from CUMULANT analysis, and mean value of the first distribution peak from CONTIN analysis of 1 µM Drp1 in presence of 0.5 mM GMPPCP, in absence (“Drp1”) or presence of 3 µM RidL (“Drp1 + RidL”) or 3 µM SidC (“Drp1 +SidC”). PDI, polydispersity index. Representative dynamic light scattering (DLS) profiles of (**B**) RidL (10 µM) or (**C**) SidC (10 µM) are shown (30 min).

**Fig. S3.**
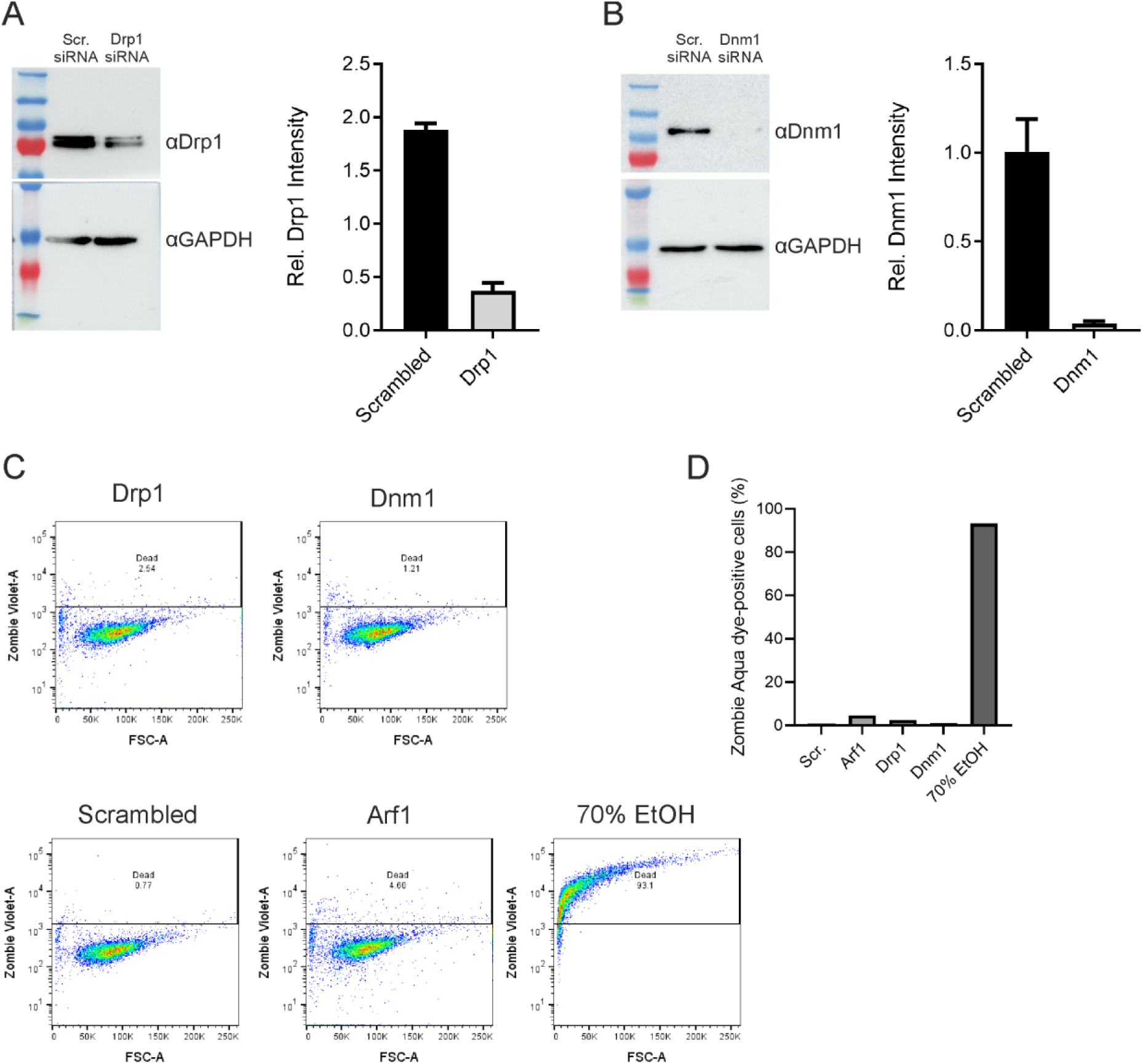
Drp1 but not Dnm1 affects intracellular replication of *L. pneumophila*. Efficiency of siRNA-mediated depletion of (**A**) Drp1 or (**B**) Dnm1 was assessed by Western blot (left panels). Quantification of the Drp1 or Dnm1 and GAPDH signal intensities of cells treated with Drp1-specific or scrambled siRNA were calculated in Image J (right panels). Graphs show the Drp1 or Dnm1 signal intensities normalized to GAPDH (n=2). (**C**, **D**) Cytotoxicity was assessed in cells transfected with siRNA or treated with 70% EtOH (30 min, 37°C) by staining with Zombie Aqua dye (1:500 dilution, 30 min). After staining, cells were fixed with 4% PFA and processed by flow cytometry with forward scatter (FSC) voltage 400, sideward scatter (SSC) voltage 250 and laser 525/50 (Vio510) voltage 380 (> 10.000 events per sample). (**C**) Scatter plots (indicated line shows the set threshold for live/dead cells) of FSC vs. Zombie Aqua dye fluorescence and (**D**) bar graph of Zombie Aqua dye-positive cells are shown.

**Fig. S4.**
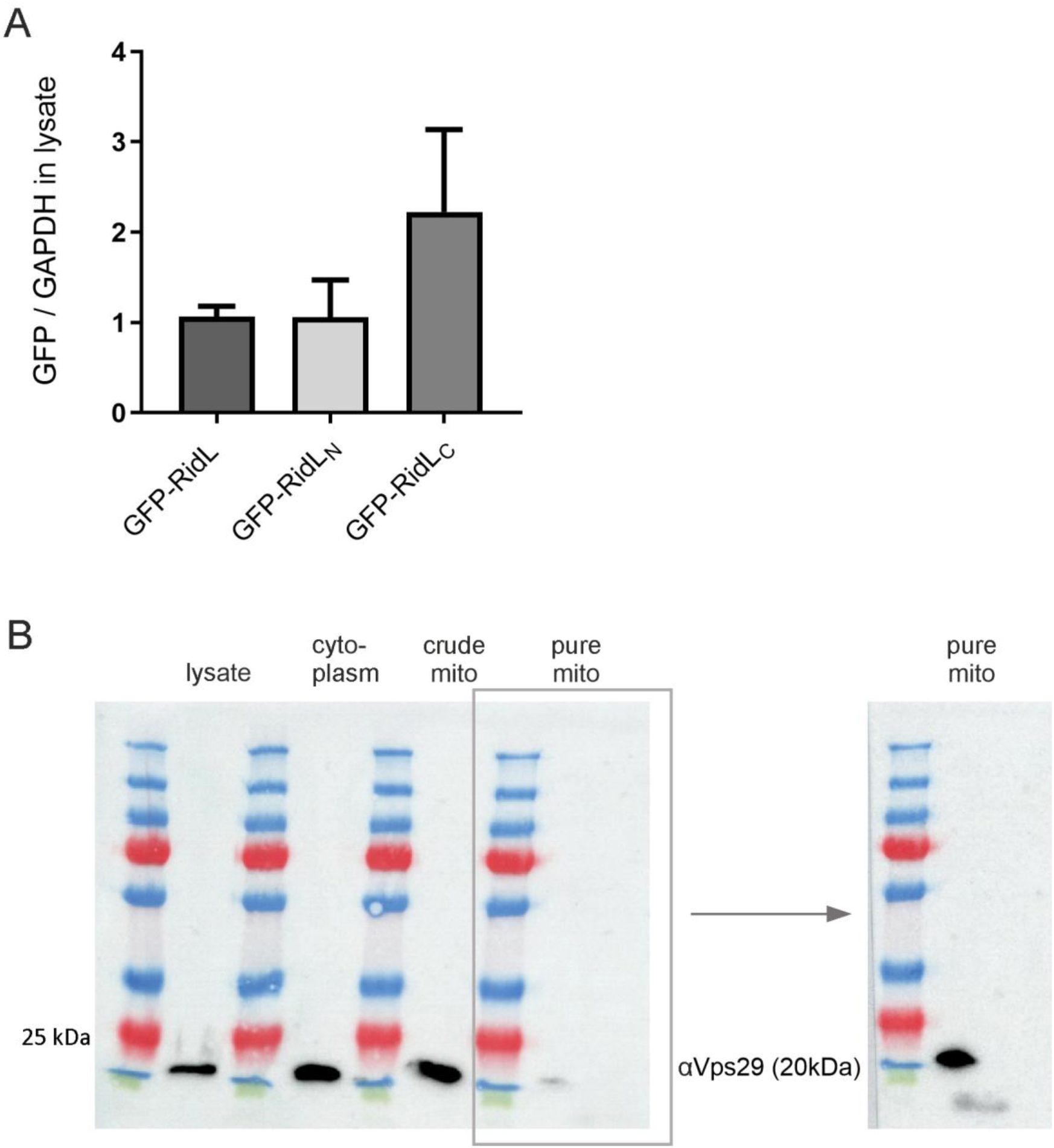
Localization of RidL and retromer. (**A**) Detection of GFP-RidL, GFP-RidL_N_ and GFP-RidL_C_ in HEK293 cell lysate. Additional information for Fig. 4C. Signal intensities of GFP-RidL, GFP-RidL_N_, GFP-RidL_C_ and GAPDH in HeLa cell lysate were assessed by Image J. Graphs show the means + SEM of relative GFP intensities (GFP/GAPDH) of three independent experiments. (**B**) Localization of the Vps29 subunit of the retromer cargo recognition subcomplex in HeLa cell lysates (lysate), cytoplasmic (cytoplasm), and crude/purified mitochondrial (crude/pure mito) fractions. Western blot with anti-VPS29 Antibody (D-1) (sc-398874, Santa Cruz Biotechnology, 1:100) and secondary antibody goat anti-mouse IgG [H+L] HRP-linked secondary antibody (Invitrogen; 1:5000). Regular (left panel) and prolonged exposure (right panel) is shown.

**Fig. S5.**
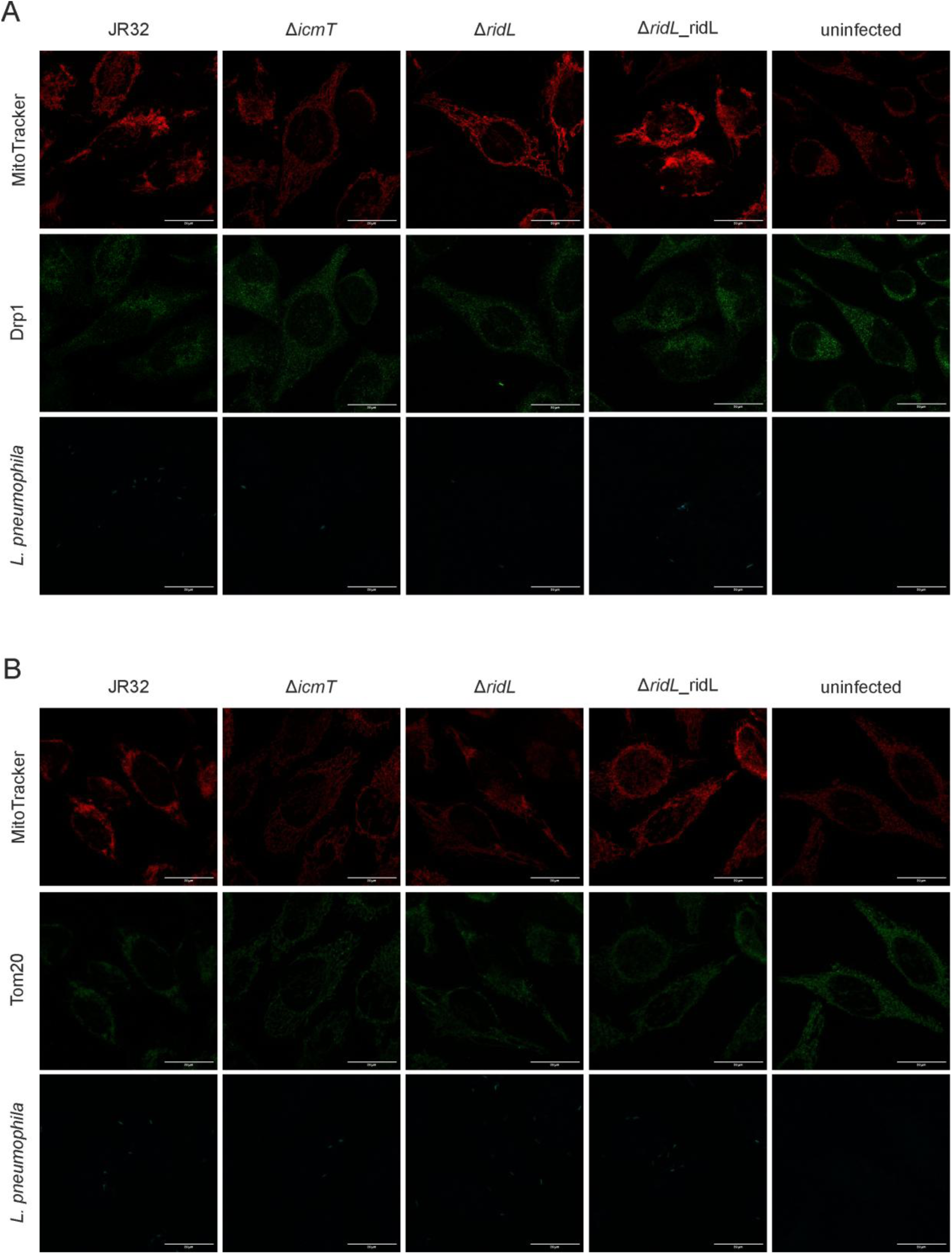
Raw microscopy images of Drp1 and Tom20 immunostaining. Images show the individual channels for MitoTracker, mCerulean (*L. pneumophila*, pNP99), and GFP of (**A**) Drp1 or (**B**) Tom20 corresponding to Fig. 4e or Fig. 4f, respectively. Original signal intensities are shown without adjustment of brightness or contrast. Scale bar = 20μm.

**Fig. S6.**
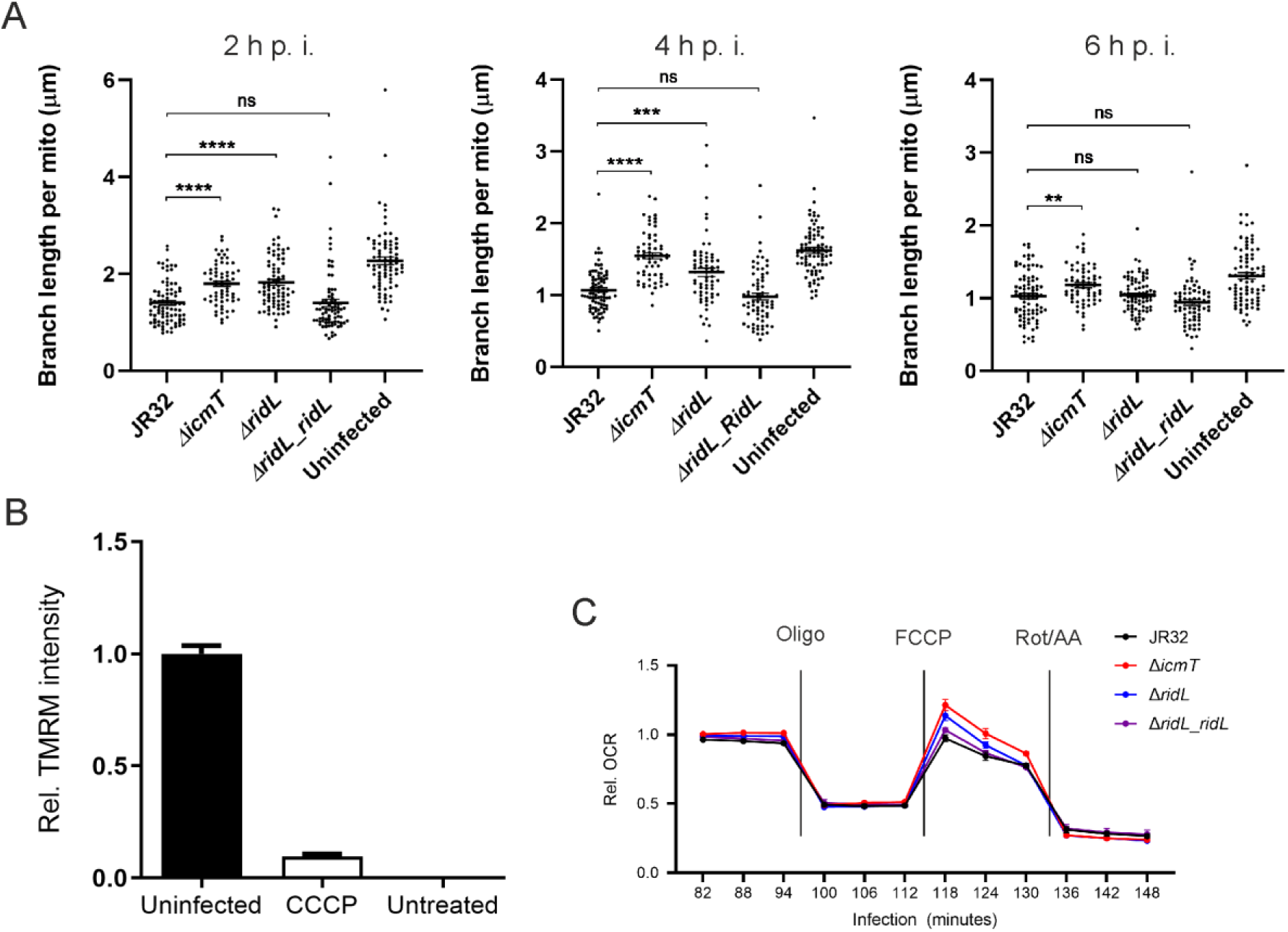
**RidL modulates mitochondrial fragmentation and function in *L. pneumophila*-infected cells**. **(A)** Mitochondrial branch lengths are quantified in infected cells (Fig. 1a). In infected cells, the average branch lengths per mitochondrion were assessed using the Mitochondria Analyzer plugin for Image J. Dots represent all analyzed cells from three independent experiments (20-30 cells each; Student’s *t*-test, **, p < 0.01; **, p < 0.001; ****, p < 0.0001). (**B**) Controls for TMRM assays shown in Fig. 5c. HeLa cells were treated with 50 μM CCCP (5 min, 37°C). Uninfected and CCCP-treated cells were stained with tetramethyl rhodamine methyl ester (TMRM; 20 nM, 30 min, 37°C), untreated cells were neither infected nor stained. TMRM intensities, reflecting the membrane potential, were assessed by flow cytometry with FSC voltage 420, SSC voltage 280 and YG610/20 voltage 400 (TMRM) (> 10.000 events per sample). Data was processed in FlowJo and normalized to the average TMRM signal intensity of uninfected cells. Graphs show means + SEM of three independent experiments (untreated mean: 0.000493). (**C**) Oxygen consumption of infected HeLa cells as shown in Fig. 5d. Oxygen consumption rates (OCR) of uninfected HeLa cells were measured 2×. HeLa cells were infected (MOI 10) with *L. pneumophila* strains JR32, Δ*icmT*, Δ*ridL* or Δ*ridL_ridL* (pNT28 or pIF009), subjected to a mitochondrial stress test, and the OCR was measured every 6 minutes in 15 cycles (3:00 mix, 0:00 wait, 3:00 measure). Oligomycin (final concentration 1 μM), FCCP (0.5 μM) and rotenone / antimycin A (0.5 μM) were injected at the indicated time points. OCR values were normalized to the average OCR of the last two time points measured before the stress test. Graphs show the OCR during infection, normalized to OCR values before infection. Means + SEM of three independent experiments.

**Fig. S7.**
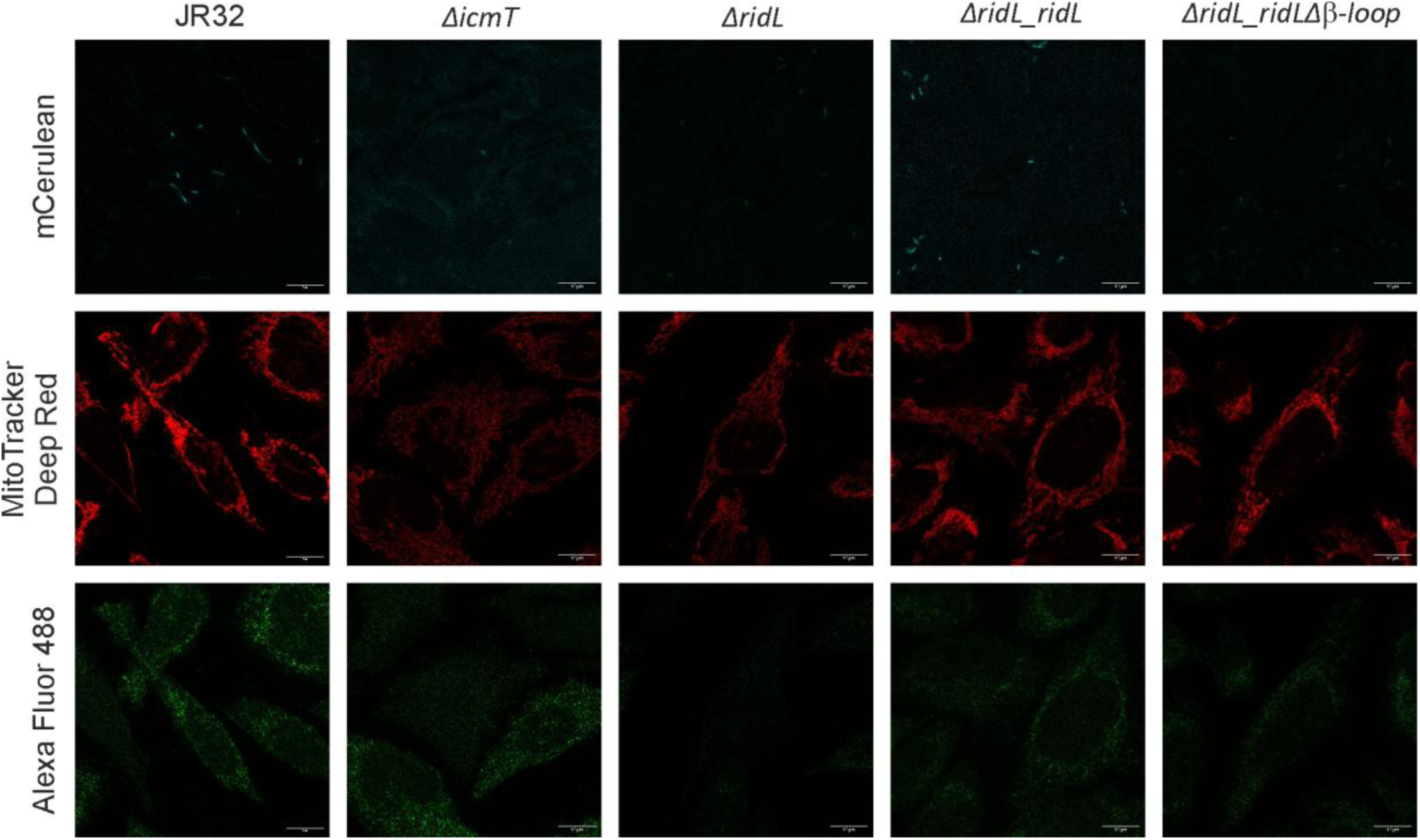
Raw microscopy images of phosph-Drp1 in *L. pneumophila*-infected. Images show the individual channels for mCerulean (*L. pneumophila*; pNP99, pKB208, or pKB209), MitoTracker Deep Red, and Alexa Fluor 488 of phospho-Drp1 (Ser616) corresponding to Fig. 6a. Signal intensities with adjustment of brightness and contrast are shown. Scale bar = 10μm.

**Table S1.** Strains and plasmids used in this study.

| Strain/plasmid | Relevant properties <sup>a</sup> | Reference |
| --- | --- | --- |
| ***S. cerevisiae*** |  |  |
| BY4741 | MATa his3Δ1 leu2Δ0 met15ΔO ura3Δ0 | Panse lab collection |
| NMY32 | MATa trp1-901 leu2-3,112 his3Δ200<br>LYS2::(lexAop)<sub>4</sub>-HIS3 ADE2::(lexAop)<sub>8</sub>-ADE2 URA3::(lexAop)<sub>8</sub>-lacZ | Dualsystems Biotech |
| ***E. coli*** |  |  |
| BL21(DE3) | *fhuA2 [lon] ompT gal (λ DE3) [dcm] ΔhsdS λ DE3 = λ sBamHIo ΔEcoRI-B int::(lacI::PlacUV5::T7 gene1) i21 Δnin5* | Novagen |
| MC1061 | *araD139 Δ(araA-leu)7697 Δ(lac)X74 galK16 galE15(GalS) λ- e14- mcrA0 relA1 rpsL150(StrR) spoT1 mcrB1 hsdR2* | <sup>2</sup> |
| TOP10 | *F- mcrA (mrr-hsdRMS-mcrBC) 80lacZ M15 lacX74 recA1 ara 139 (ara-leu)7697 galU galK rpsL (StrR) endA1 nupG* | Invitrogen |
| ***L. pneumophila*** |  |  |
| CR06 (ΔridL) | *L. pneumophila* JR32 *ridL::Kan<sup>R</sup>* | <sup>3</sup> |
| GS3011 (ΔicmT) | *L. pneumophila* JR32 *icmT3011::Kan<sup>R</sup>* | <sup>4</sup> |
| JR32 | Virulent *L. pneumophila* sg 1 strain<br>Philadelphia | <sup>5</sup> |
| **Plasmids** |  |  |
| pAct2.2-large T | Control | Panse lab collection |
| pBXNH3 | N-terminal 3C protease cleavable His<sub>10</sub>-tag, pBAD, Amp<sup>R</sup> | <sup>6</sup> |
| mCh-Drp1 | Drp1 (human) | Addgene #49152, <sup>7</sup> |
| pET21-Drp1 | Drp1 (murine) | Addgene #72929, <sup>8</sup> |
| pEV001 | pET21-Drp1 (human) | This work |
| pEGFP-N1-dynamin | Dynamin-eGFP | Addgene #120313, <sup>9</sup> |
| pIF009 | pMMB207-C-RBS-*gfp*-P<sub>ridL</sub>-RidL<br>(constitutive GFP) | <sup>3</sup> |
| pKA001 | pLexA-RidL | This work |
| pKA004 | pAct2.2-Vps1 (yeast) | This work |
| pKA009 | pAct2.2-Drp1 (human) | This work |
| pKA103 | pAct2.2-Dnm1 (human) | This work |
| pKA011 | pAct2.2-Dnm2 (human) | This work |
| pKA104 | pAct2.2-Dnm3 (human) | This work |
| pKA014 | pAct2.2-DymA (*D. discoideum*) | This work |
| pKA015 | pAct2.2-DymB (*D. discoideum*) | This work |
| pKA067 | pRS315_Nop1pr_GFP11-mCherry-RidL | This work |
| pKA072 | pRS316_Nop1pr_GFP1-10-Vps1 | This work |
| pKA100 | pRS316_Nop1pr_GFP1-10-Drp1 | This work |
| pKA097 | pRS316_Nop1pr_GFP1-10-Dnm1 | This work |
| pKA098 | pRS316_Nop1pr_GFP1-10-Dnm2 | This work |
| pKA099 | pRS316_Nop1pr_GFP1-10-Dnm3 | This work |
| pKA105 | pGEM-T Easy_NdeI-DymB-BamHI | This work |
| pKA114 | pRS316_Nop1pr_GFP1-10-DymA | This work |
| pKA115 | pRS316_Nop1pr_GFP1-10-DymB | This work |
| pKA117 | pGEM-T-Easy_DymB | This work |
| pKA118 | pRS316_Nop1pr_GFP1-10-AgeI | This work |
| pKA080 | YEpl351tet-Drp1 | This work |
| pKA173 | pET21-Drp1 (human) | This work |
| pKB002 | pINIT-RidL | 10 |
| pKB139 | pBXNH3_RidL<sub>1-258</sub> | This work |
| pKB198 | pMMB207C-GFP-P<sub>ridL</sub>-RidL$\Delta\beta$-loop | This work |
| pKB208 | pMMB207C-mCerulean-P<sub>ridL</sub>-RidL | This work |
| pKB209 | pMMB207C-mCerulean-P<sub>ridL</sub>-RidL$\Delta\beta$-loop | This work |
| pKB241 | pINIT-Dnm2 (PCR template) | This work |
| pKB242 | pINIT-Vps1 (PCR template) | This work |
| pKB245 | pEGFP-N1_RidL<sup>opt</sup> (PCR template) | This work |
| pKB248 | pEGFP-C1_RidL<sup>opt</sup>-STOP (eGFP-RidL) | This work |
| pKB249 | pEGFP-C1_RidL<sup>opt</sup><sub>9-258</sub>-STOP | This work |
| pKB250 | pEGFP-C1_RidL<sup>opt</sup><sub>259-1167</sub>-STOP | This work |
| pKB306 | pBXC3GH-RidL<sub>437-1100</sub> | This work |
| pLexA_Lamin C | Control | Panase lab collection |
| pNT28 | pMMB207-C, $\Delta lacI^q$ (constitutive *gfp*),<br>Cam<sup>R</sup> | 11 |
| pNP99 | pMMB207-C, $\Delta lacI^q$ (constitutive<br>mCerulean), Cam<sup>R</sup> | 12 |
| pSJ1321 | pRS315-NOP1pr-GFP<sub>11</sub>-mCherry-PUS1 | Addgene #86413, <sup>13</sup> |
| pSJ2039 | pRS316-NOP1pr-GFP<sub>1-10</sub>-SCS2TM | Addgene #86418, <sup>13</sup> |

<sup>a</sup> Abbreviations: Amp, ampicillin; Cam, chloramphenicol; Kan, kanamycin.

**Table S2.**
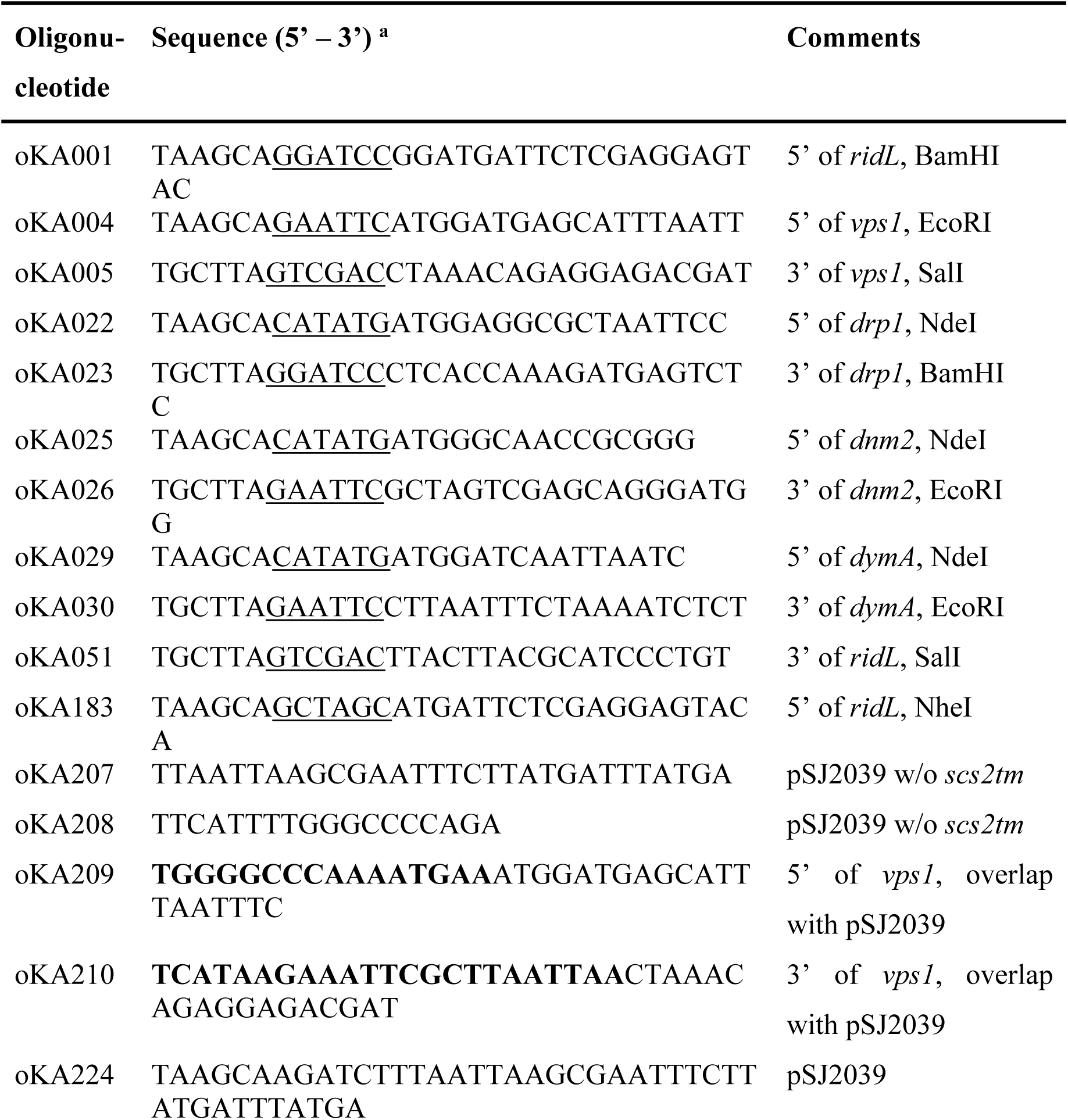
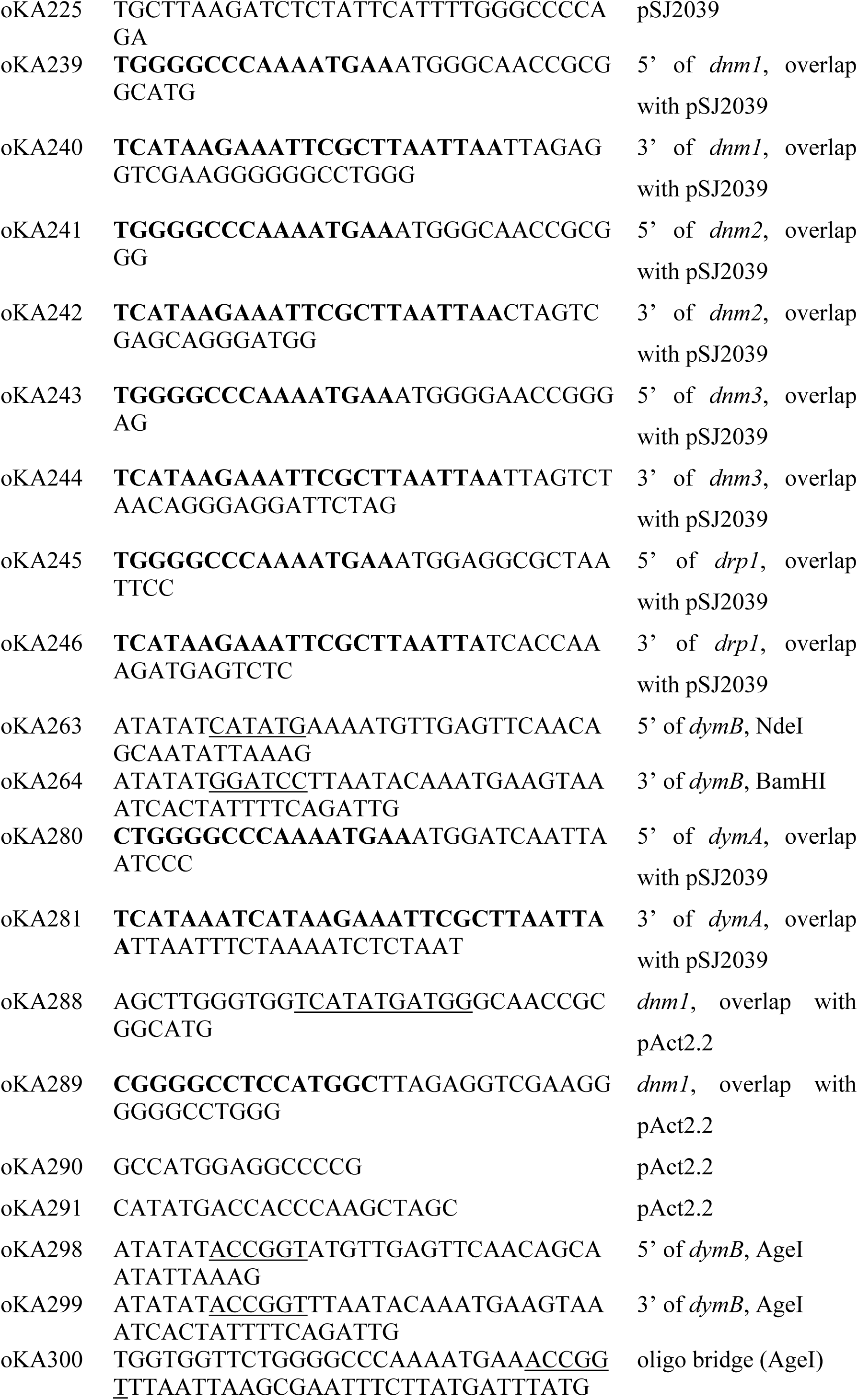
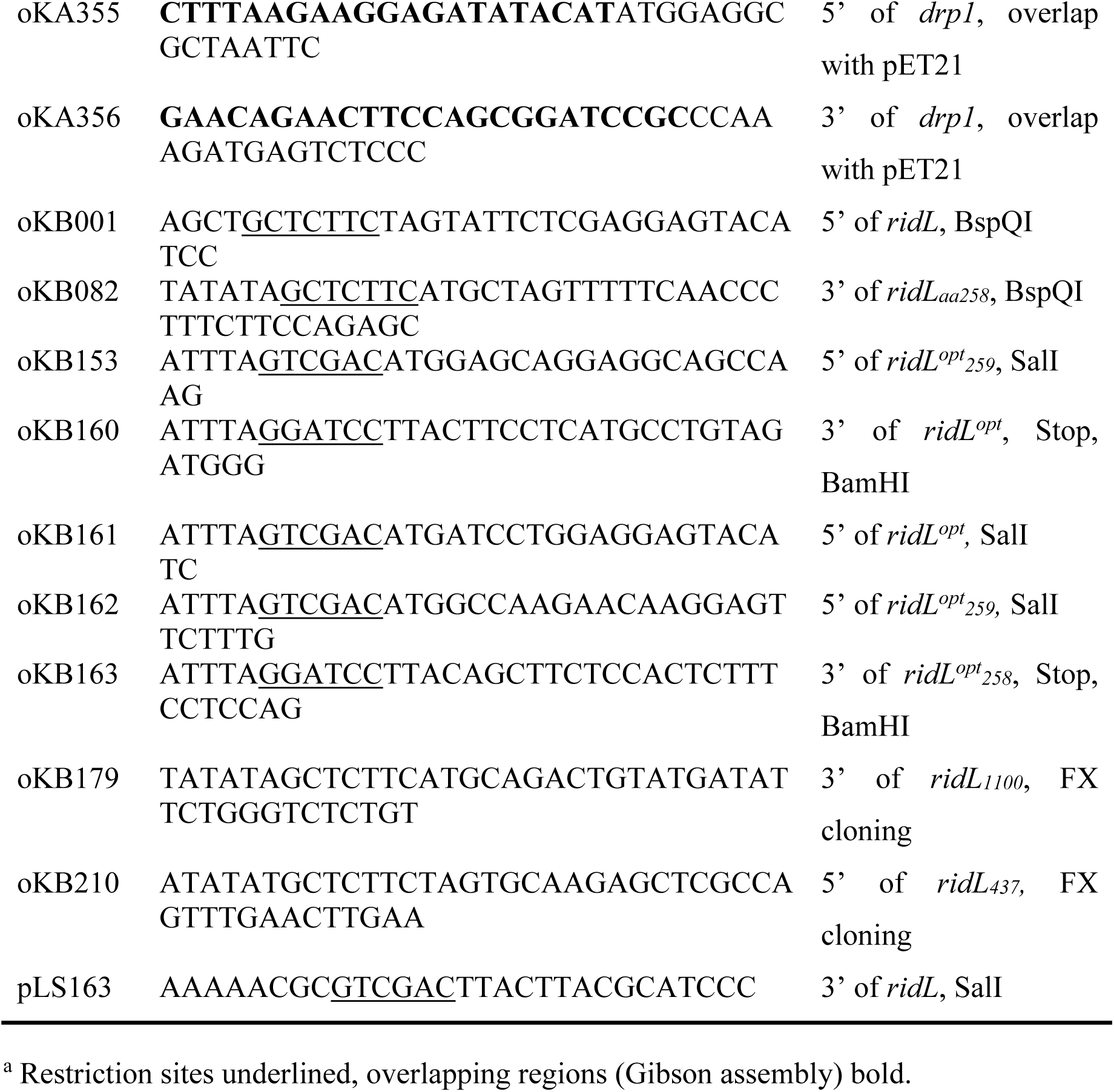
Oligonucleotides used in this study.

